## Supporting Information for "Genome evolution is surprisingly predictable after initial hybridization"

*Supporting Information 1. Description of the X. cortezi genome assembly*

In addition to annotating protein coding genes in the *X. cortezi* genome (see main text), we annotated repetitive elements. We used RepeatMasker with a custom library of poecilid transposable elements (see Methods), which resulted in the identification of 278 Mb of repetitive sequences, largely DNA/TcMar, DNA/hAT, and LINE/L2 elements (Table S1). The most differentiated transposable element classes between the *X. birchmanni* and *X. cortezi* assemblies were LTR/ERV-Foamy elements which were twice as common in the *X. birchmanni* assembly (but only make up 2.8 Mb of the 723 Mb assembly; Table S1).

We were also interested in rearrangements that might differentiate *X. birchmanni* and *X. cortezi.* Using approximate alignments generated with minimap2 [1], we evaluated synteny across the two assemblies, and identified inverted and translocated rearrangements between the two species that exceeded 100 kb in length. We found that the two genomes were largely syntenic, with evidence of only a dozen rearrangements of this size. We found evidence for three translocations, which we treat with caution in the absence of Hi-C data for *X. cortezi*, and nine inversions ranging in size from 218 kb to 6.7 Mb (Table S2). These inversions were concentrated on chromosomes 8 and 17 (six out of nine of the inversions). We note that with one sampled individual per species, we cannot distinguish between segregating and fixed structural rearrangements, but given low genetic diversity within species relative to divergence between species ($\pi \sim 0.1\%$ versus $D_{xy} \sim 0.6\%$), we predict that the majority of these rearrangements will represent fixed differences.

*Supporting Information 2. Discussion of PCA and relatedness analyses of high-coverage hybrid individuals*

In the main text, we describe a series of analyses intended to evaluate genetic variation and relatedness between the two *X. birchmanni* x *X. cortezi* hybrid populations. In particular, we were interested in evaluating any signals in the Santa Cruz and Chapulhuacanito populations that might indicate that these hybridization events were not truly independent. Although the results of these analyses support the inference that the hybrid populations are independent (see Results), they point to some other unexpected signals that we discuss in more detail here.

We identified regions that were homozygous for *X. cortezi* ancestry in all six of our high coverage hybrid individuals (three from Santa Cruz and three from Chapulhuacanito), and extracted SNPs that fell in those ancestry tracts in hybrids and pure *X. cortezi* individuals (Fig. 2C). Separately, we identified regions that were homozygous for *X. birchmanni* ancestry in all six hybrid individuals and extracted SNPs that fell in those ancestry tracts in hybrids and pure *X. birchmanni* individuals. For this analysis, we included *X. birchmanni* from an allopatric source population (Coacuilco), and pure sympatric *X. birchmanni* individuals (Fig. 2D). Our expectation for these analyses was that ancestry tracts from the two hybrid populations would separate from each other, which we observe, but that they would cluster more closely with ancestry tracts from the *X. birchmanni* individuals with which they are sympatric, as these individuals presumably are closely related to the individuals that contributed *X. birchmanni* ancestry in the hybridization event. However, we instead see that the *X. birchmanni* individuals collected from Santa Cruz and Chapulhuacanito cluster closely with each other and with the allopatric *X. birchmanni* population (Fig. 2D). Notably, we observe a similar result in GCTA analysis of the genetic relatedness matrix, with some *X. birchmanni* individuals in different populations identified as more related than average (see Methods, Fig. S21).

We are unsure of the driving forces behind this pattern but discuss several possibilities here. First, it is possible that errors in inference of the *X. birchmanni* ancestry tracts (which are more difficult to accurate delineate since they are smaller, see Supporting Information 7), introduce divergent SNPs into the PCA analysis of *X. birchmanni* ancestry tracts and result in more separation between hybrids and the source *X. birchmanni* populations than expected a priori. Second, a small number of *X. birchmanni* individuals could have contributed to the initial hybrid population, resulting in genetic drift between *X. birchmanni* tracts in hybrids and sympatric *X. birchmanni* individuals. Other explanations could include that the population history of *X. birchmanni* has driven higher than expected genetic similarity between individuals from geographically distinct populations. PSMC analyses suggest that *X. birchmanni* experienced a recent bottleneck within the last ~1,000 generations [2]. Finally, dispersal distances for *X. birchmanni* could greatly exceed what has been previously assumed for this small bodied species. These explanations are also consistent with positive values of genetic relatedness detected for *X. birchmanni* in some cross-population comparisons in GCTA analyses (Fig. S21).

*Supporting Information 3. Comparisons of the X. birchmanni and X. cortezi recombination map*

Past work on swordtails and the larger group to which they belong (percomorph fish) has indicated that all species studied to date carry a truncated version of PRDM9, the regulator of recombination hotspots in mammals. This version lacks the KRAB and SSXRD protein domain and is thus expected to be inactive, since these domains are essential to its function in mammals [3,4]. Consistent with this inference, both linkage disequilibrium based maps (in *X. birchmanni*) and crossover based maps (in *X. birchmanni* x *X. malinche* hybrids) indicate that recombination events largely localize to promoter-like elements including transcriptional start sites, CpG islands, and H3K4me3 peaks detected in testis ChIP-seq [2,3].

Given that species that lack PRMD9 and localize recombination events to promoter-like elements tend to have slowly evolving recombination maps, even over millions of generations [5,6], we expected that *X. birchmanni* and *X. cortezi* would likely have nearly-identical recombination maps. However, we chose to investigate this directly by building a recombination map for *X. cortezi*, to complement the linkage-disequilibrium based map we had previously generated for *X. birchmanni* [2]. We also updated the *X. birchmanni* map with our most recent genome assembly (see Methods), which is much more complete than previous assemblies (NCBI submission ID: JAXBVF000000000).

Given that we sequenced fewer individuals in *X. cortezi* and lack access to pedigree with which to estimate mendelian errors and improve phasing (see Methods), we expect this map to be lower resolution and to have a higher error rate than the *X. birchmanni* map. Thus, we compared the two maps qualitatively to confirm that patterns of local recombination rate variation are generally concordant. Since the maps were generated in different coordinate space, we lifted over the *X. cortezi* map to *X. birchmanni* coordinate space using haltools [7]. We then compared the average ρ/bp across species in windows of varying size and found that the *X. birchmanni* and *X. cortezi* maps were strongly correlated (Fig. S5).

In previous work, we performed simulations to ask about how accurately we might expect LD-based methods to reflect the true recombination rate [2]. In these simulations we found that, as expected, inferred LD-based maps were strongly but imperfectly correlated with the true recombination map [2], with a median Spearman's correlation across simulations between the true and inferred maps of ~0.65 (range: 0.59-0.71 in an analysis of 50 kb windows). In our comparisons of *X. birchmanni* and *X. cortezi* over the same spatial scale, we find that the Spearman's correlation between the two maps is 0.55 (Fig. S5), slightly lower than expected if the two maps were in fact identical. Given differences in population history (see next paragraph), expected differences in error rates between the maps, and chromosome architecture (Fig. S15), we consider this evidence that the *X. birchmanni* and *X. cortezi* maps are extremely similar.

We note that the median ρ/bp in *X. birchmanni* and *X. cortezi* is inferred to be different from our analyses. In *X. birchmanni*, the median ρ/bp is 0.00076 whereas in *X. cortezi* it is 0.0026. Since ρ reflects the composite value of 4*Ner*, we asked whether differences in *Ne* of *X. birchmanni* and *X. cortezi* could in part explain this difference. We reanalyzed PSMC results for individuals from the Río Huichihuyan populations of *X. cortezi* and the Coacuilco population of *X. birchmanni* from previous work [8] and asked about the inferred *Ne* of *X. birchmanni* and *X. cortezi* over the past 100,000 generations. We assumed the same generation time and mutation rate for these two species (following [8]). Using this approach, we estimated that *X. birchmanni* has had a long-term effective population size of approximately 33k over the last 100,000 generations, whereas *X. cortezi* has had an *Ne* of approximately 66k over the same time period. This suggests that the differences in estimated ρ/bp in *X. cortezi* are in part attributable to differences in population size, although differences in error rates across the two maps may also contribute.

*Supporting Information 4. Simulations to explore observed cross-population correlations*

We observe strikingly strong correlations in local ancestry across the Santa Cruz and Chapulhuacanito populations. Since these two populations do not have a shared demographic history, cross-population correlations are likely generated in part by shared sources of selection across the two populations. Another possible source of shared patterns of local ancestry could be shared sources of error in local ancestry inference (see Supporting Information 5-7).

Under neutrality, cross-population correlations in local ancestry are unexpected (Fig. S19). To explore whether selection could in principle generate the patterns of cross-population ancestry correlations we observe, we performed simple simulations, without attempting to directly infer the architecture of selection on *X. birchmanni* x *X. cortezi* hybrids or to simulate the full range of possible scenarios of selection on hybrids [9]. Our previous work identified 81 minor parent ancestry deserts in the Santa Cruz population [10], pointing to a reasonable starting number for simulations of loci under selection in *X. birchmanni* x *X. cortezi* hybrids.

We used the admixem program [11] to simulate admixture and selection on hybrids. We modeled 24 chromosomes, equivalent in length to the *X. birchmanni* chromosomes, and used the empirical recombination map from *X. birchmanni* to specify crossover probabilities. For each simulation, we drew other parameters, such as generations since initial admixture and initial admixture proportion, from the posterior distributions of ABC demographic inference simulations for Santa Cruz and Chapulhuacanito respectively. To model selection, we simulated recessive Dobzhansky-Muller hybrid incompatibilities. We simulated 40 incompatibilities (involving 80 loci). For each simulation, we randomly drew the chromosome and position of each locus in an incompatibility. Based on the results of initial simulations, we drew the selection coefficient from a random exponential distribution with a mean of 0.6. This distribution was truncated such that values could not exceed 1.

Because the ABC approach that we used to infer demographic history lacked selection, we found that using the posterior distribution of inferred initial admixture proportions in simulations with selection resulted in nearly complete purging of simulated *X. birchmanni* ancestry. This observation is in line with what other researchers have reported based on both empirical and theoretical results [12–14]. As a result, we modified initial admixture proportions in the simulations by 20% and found that in practice this results in final admixture proportions that overlapped with those observed in the empirical dataset for both simulated populations (Fig. S18 compared to Fig. 1).

We compared 50 replicate simulations of two hybrid populations. These populations differed in the simulated demographic history, but experienced selection on the same pairs of hybrid incompatibility loci. We summarized the results of these simulations by calculating the average ancestry in 250 kb windows across the genome. We examined correlations in local ancestry across the two replicate populations for each simulation as we had for the real data. We found that correlations in local ancestry in simulated data overlapped those observed in the real data under this scenario of extremely strong selection (Fig. S19).

We repeated these simulations modeling partial dominance of incompatibility loci, by drawing the $h$ parameter from a random uniform distribution from 0-0.5. Given that setting $h > 0$ will decrease the average fitness of hybrids, we changed the average selection coefficient for this set of simulations to 0.4, but otherwise performed the simulations as described above. We again found that these simulations under this scenario resulted in correlations in ancestry that overlapped those observed in the real data (Fig. S19).

Together, these results indicate that in principle very strong selection on hybrids can drive the level of cross-population correlations in ancestry that we observe in comparisons between the Santa Cruz and Chapulhuacanito populations.

*Supporting Information 5. Simulations of expected accuracy in local ancestry inference*

To evaluate the expected accuracy of our local ancestry inference approach, we used the program *mixnmatch*. *mixnmatch* is a pipeline designed by our lab [15] that simulates divergence between the parental lineages and admixture in hybrid populations, performs local ancestry inference, and evaluates the accuracy of that inference relative to the true simulated ancestry. We performed these simulations matching the inferred demographic history of the Santa Cruz and Chapulhuacanito populations from ABCreg analysis. To simplify these simulations, we used the MAP estimates for each demographic parameter (Fig. 1; Fig. S1) and set the hybrid population size to 5,000. We used *mixnmatch* to simulate haplotypes for 50 diploid admixed individuals from each population and generate reads from each individual. We then ran *ancestryinfer* as we had on the real data and compared inferred ancestry at a posterior probability threshold of 0.9 to true ancestry in each simulated individual. As expected based on previous results [8,15], we found that error rates in local ancestry inference are expected to be very low (Fig. S11).

We also evaluated how shared clusters of errors might impact the correlations in ancestry inferred across populations. To do so, we performed admix'em simulations similar to those described above (Supporting Information 4), except that we did not implement selection. Briefly, we simulated 24 chromosomes matching the length of the *Xiphophorus* chromosomes, tracked ancestry at 1,000 markers per chromosome, and drew simulation parameters for each population from the posterior distributions of demographic parameters produced by ABCreg. We performed 20 replicate simulations.

After the simulations had finished running, we then artificially added clusters of errors to the data. Our estimated error rate from pure parentals and artificial hybrids is ~0.1% (see Methods). However, to be conservative, we chose to simulate a high error rate of 5%. We reasoned that errors that occurred in clusters would have a larger impact on cross-population correlations than sporadic errors. As such, we randomly sampled 400 markers, and generated error tracts that continued for 3 markers (approximately 75 kb in our simulations). In both simulated populations, we converted the ancestry calls within these tracts to homozygous *X. birchmanni* ancestry in all individuals (i.e. minor parent ancestry). We then calculated average ancestry in 250 kb windows as we had in the real data, and calculated the Spearman's correlation in ancestry across the paired simulations. We repeated this procedure 20 times.

Based on the results of these high-error simulations, we found that cross-population ancestry correlations did not approach those observed in the real data (median Spearman's $\rho$ = 0.32; range 0.21-0.47). Even in this conservative scenario of very high error rates and identical error profiles across all individuals, we are not able to replicate the patterns seen in our data with error alone. Together these results make us confident that the strong cross-correlations in ancestry we observe across *X. birchmanni* x *X. cortezi* populations are at least in part attributable to selection rather than errors in ancestry inference.

*Supporting Information 6. Evaluating other factors that could impact correlations in ancestry across populations*

Given the extraordinarily high correlations in local ancestry that we observe between the Santa Cruz and Chapulhuacanito populations, we wanted to rigorously evaluate potential technical factors that could contribute to this pattern and be mistaken as biological signal. In addition to the analyses described in the main text (see Methods) and simulation-based methods described above, we pursued a number of additional analyses where we filtered our data using different criteria and re-evaluated correlations in local ancestry across populations.

While they are rare, we do detect errors in local ancestry inference in our analyses of known crosses (see Methods). To further investigate potential effects of error rates in the real data, we identified all likely errors genome-wide in early generation hybrids. We defined errors as instances where we observe a switch between ancestry states that reverts to the original ancestry state within 100 kb (Fig. S23). We then excluded these regions from our data from Santa Cruz and Chapulhuacanito and asked about the observed cross-correlations in ancestry in 100 kb windows. Because excluding windows with errors in any early generation hybrid results in a reduced number of windows for analysis, we generated a comparison dataset of 1,000 size-matched datasets where a matched number of windows was randomly sampled from the non-filtered data. We next compared the Spearman's correlation in the filtered and 1,000 size-matched unfiltered datasets. The observed correlation in ancestry between the two hybrid populations in the filtered dataset was not significantly lower than that observed in size-matched unfiltered datasets (filtered = 0.76, versus average size matched = 0.78; p-value by simulation = 0.16).

We next explored the effects of removing regions of the genome where errors might be expected to be more common, but where we lacked direct evidence that errors had occurred. First, we removed all ancestry informative markers that overlapped with annotated repeats, where errors may be more common, and found that the correlations in local ancestry were unchanged (Spearman's ρ in 250 kb windows - 0.82). We also removed windows within 1 Mb of the end of the assembled chromosome, given that error rates may be higher in these regions since some chromosomes contain telomeric sequences (Spearman's ρ in 250 kb windows - 0.84; Fig. S15). In addition to the thinning approaches we used to evaluate the impacts of power in the main text (see Methods), we further removed all windows where the number of ancestry informative sites in the window fell in the lowest 5% or 10% genome-wide and re-evaluated correlations between populations (Spearman's ρ 0.82-0.83 in 250 kb windows across comparisons). We performed similar analyses excluding windows where we have the highest power to infer ancestry (upper 5 or 10% of ancestry informative site density) and again saw that correlations in ancestry between the two populations remained strong (Spearman's ρ 0.79-0.81 in 250 kb windows across comparisons). We also evaluated combinations of these analyses (i.e. removing repetitive windows, windows within 1 Mb of chromosome edges, and low-power windows). In no case did these modifications substantially change the observed cross-population correlations.

In summary, our analyses in the main text and those described above indicated that errors and variation in power to infer ancestry were very unlikely to generate the cross-population correlations we observe in the real data. However, we have a clear expectation that ancestry correlations across distinct hybrid populations should be substantially lower than those observed when analyzing ancestry covariance within the same population. Thus, we wanted to evaluate the range of correlations expected from subsampling individuals from the same population (in addition to the cross-year and within-drainage comparisons discussed in the main text) and to compare these with our cross-population results. We generated replicated datasets where we randomly subsampled 20 individuals from the same population and collection year and calculated average ancestry along the genome. Across these subsampled replicates, we consistently observed correlations in local ancestry in 250 kb windows of ≥0.9 in subsampled replicates from both Santa Cruz and from Chapulhuacanito. Reassuringly, these correlations from subsampling the same population greatly exceed the correlations observed when comparing ancestry across the two populations.

Overall, our results are not consistent with technical factors driving cross-population correlations (see also Supporting Information 5) and instead point to biological factors, such as shared sources of selection, driving the genome-wide correlations in ancestry that we observe in *X. birchmanni* x *X. cortezi* hybrids.

*Supporting Information 7. Additional analyses and discussion of spatial patterns from the Discrete Wavelet Transform analysis*

We performed wavelet analyses of correlations using two different interpolation resolutions of the ancestry estimate. The ancestry statistic is an estimate of the minor parent allele frequency within a diploid individual, calculated as a weighted average of marginal posterior probabilities of two genotypes:

$$f_A = P(AA) + \frac{1}{2} P(Aa)$$

We subsequently average these interpolated measures across individuals to give an estimate of the population admixture proportion. In one case, we interpolate ancestry to a resolution of 32 kb. This corresponds roughly to the average tract length of minor parent ancestry in the Santa Cruz population. We separately interpolate to a 1 kb grid, corresponding roughly to the density of ancestry informative markers.

Our analyses of cross-population ancestry correlations revealed strong correlations persisting at fine genomic scales (Fig. S13). We interpret these correlations with caution, as they could be in part driven by correlated fine-scale error in the ancestry inference procedure (see details below). Moreover, we also expect that since the majority of ancestry 'tracts,' defined on the basis of high-confidence transitions between ancestry states, are greater than 32 kb in both *X. birchmanni* x *X. cortezi* hybrid populations, local ancestry inference will be less reliable at these fine scales (see also below). That said, some of these fine scale correlations may indeed be capturing real biological signals. This is because the wavelet analysis directly uses the posterior probability of each ancestry state. As a result, even short ancestry tracts that are not called with high confidence could generate spatial variation in marginal posterior probabilities that correlates with true ancestry states. Furthermore, we might expect that a bias towards detection of minor parent ancestry would be stronger in lower recombination regions. However, if this were true, we would expect to see negative correlations between minor parent ancestry and recombination at these scales, which we do not (Fig. S12). Finally, we note that the pattern we observe is consistent with expectations that hybrid populations formed between more deeply divergent species will have a higher density of selected sites. Fine-scale correlations in the analysis of *X. birchmanni* x *X. cortezi* populations are substantially elevated compared to correlations at fine-scales in analysis of *X. birchmanni* x *X. malinche* populations (Fig. S13). This pattern suggests that the repeatability we observe at small spatial scales in *X. birchmanni* x *X. cortezi* populations may be driven by stronger selection on hybrids in this more divergent cross.

While we expect our local ancestry inference approach to be highly accurate overall, there are several reasons to be cautious about patterns detected at the finest spatial scales. Past simulation studies have indicated that *ancestryinfer* has a much higher error rate in shorter ancestry tracts [15]. Shorter ancestry tracts will contain fewer ancestry informative sites with which to infer local ancestry. Moreover, because of concerns about inducing correlations between recombination rate and minor parent ancestry [15], *ancestryinfer* uses a uniform recombination prior, which may further increase the difficulty of detecting short ancestry tracts.

To evaluate expected error rate in short ancestry tracts directly, we used *mixnmatch* simulations matching the demographic history of the Santa Cruz population. We expect that simulations of Santa Cruz will result in a higher estimates of error in shorter ancestry tracts than simulations of Chapulhuacanito. This is because Santa Cruz has an older estimated admixture time (Fig. 1) and a more skewed admixture proportion towards *X. cortezi*, which will both contribute to shorter minor parent ancestry tracts [16]. Using *mixnmatch* simulations, we evaluated accuracy in simulations of 50 individuals as a function of ancestry tract length. While genome-wide error rates per ancestry informative site are estimated to be <0.4% (see *Supporting Information 5*), in ancestry tracts ≤20 kb in length, the error rate approaches 2%, and in tracts ≤10 kb in length, the error rates jump to nearly 5%. A closer examination of these errors indicates that they are almost exclusively caused by ancestry switches to homozygous major parent ancestry within heterozygous ancestry tracts (94% of errors). As a result, we interpret results of wavelet analyses at small spatial scales with caution and refrain from discussing them in the main text.

*Supporting Information 8. Results of power simulations evaluating our ability to identify shared regions under selection*

Given that we identify ~40 shared minor parent ancestry deserts between the two *X. birchmanni* x *X. cortezi* populations in the real data, we were interested in exploring our power to identify shared ancestry deserts at a range of selection coefficients. We used admix'em simulations as described above (Supporting Information 4) to simulate selection against minor parent ancestry in two populations at a range of strengths ($s$=0.01 – 0.1, with $h$=0.5). As before, for each replicate simulation, we drew from the posterior distribution of ABCreg analysis to set demographic parameters for paired simulations modeling admixture in Santa Cruz and Chapulhuacanito. We randomly determined the chromosome and position of the selected site in each simulations. We performed 100 replicate simulations per selection coefficient.

For each simulation, we asked whether an ancestry desert was detected in Santa Cruz, Chapulhuacanito, or both, as we had for the real data (see Methods). If this region overlapped with the true site under selection, we treated it as a true positive, and if it occurred on a chromosome where selection was not occurring, we treated it as a false positive. We used the proportion of time the desert overlapped with the true site under selection as an estimate of our power to detect deserts at a given selection coefficient.

Overall, we found that we had excellent power to detect stronger selection (s=0.1) in both Santa Cruz and Chapulhuacanito, and good power to detect more moderate selection (Fig. S24; >50% of selected sites with s=0.025 and ~70% of selected sites with s=0.05). We also found that the false positive rate was low (with an average of 2-4 false positives detected per simulation).

These simulation results suggest that we have excellent power to identify shared ancestry deserts in Santa Cruz and Chapulhuacanito, even when selection is modest. Moreover, the expected false positive rate is quite low. This indicates that the shared ancestry deserts identified in our empirical data are likely true sites under selection in *X. birchmanni* x *X. cortezi* hybrids, providing exciting candidates for further work exploring hybrid incompatibilities and the architecture of reproductive isolation between these two species.

*Supporting Information 9. Enrichment analysis of genes occurring in shared islands*

Since regions of high minor parent ancestry in both populations may reflect regions that have adaptively introgressed from *X. birchmanni*, we were interested in exploring whether any functional classes might be over-represented in shared minor parent islands using a gene ontology based approach. We lifted over the coordinates of shared minor parent islands identified in both Santa Cruz and Chapulhuacanito to the *X. maculatus* 5.0 (GCA_002775205.2) assembly using haltools. We then used bedtools to identify the Ensembl gene ids of the genes that fell within these islands.

*X. maculatus* Ensembl IDs were matched with GO term IDs using the R packages 'biomaRt' [17] and 'GOstats' [18]. This set of genes was used as the reference gene set. To identify GO terms with overrepresentation within the high *X. birchmanni* ancestry genes in the minor parent islands, we used a hypergeometric test implemented through the R package 'GSEABase' [19] to statistically compare observed gene counts with expected gene counts per GO term, given the reference gene set. We ran this analysis for the biological pathway, molecular function, and cellular component categories, and used a hypergeometric test p-value cutoff of 0.05 to pull significantly enriched GO terms.

Because genes with similar function sometimes colocalize in the genome, we wanted to build null expectations for gene otology enrichment that might be expected by chance. To do so, we generated null islands. For each island, we simulated a "null island" by randomly selected a start position in the genome and setting the stop position to the start position plus the length of the islands. We repeated this until we had a complete dataset of null islands and generated a total of 10 null datasets. Next, we repeated the gene otology enrichment analysis we had applied to the real data and evaluated how many and which categories were enriched in our null datasets.

Although we identified significantly enriched gene otology categories (Table S14), we avoid interpreting these results based on the results of the null datasets discussed above. Specifically, we find that neither the number of gene otology categories identified in the real analysis, nor the specific categories identified are unusual when compared to the null datasets.

*Supporting Information 10. Other drivers of variation in minor parent ancestry*

We were interested in annotating genes involved in protein complexes since past work in swordtails has highlighted that mismatch in ancestry in protein complexes can have a substantial impact on hybrid survival [20]. We used a large database of curated protein complexes annotated in humans [21] and identified reciprocal best blast hits to this database. Although swordtails are distantly related to humans, this represents the most complete dataset of protein complexes available for any species.

We used the program orthologr and the command blast_rec [22] to identify the reciprocal best blast hit for all protein coding genes in the human genome. We used the hg38 assembly (Homo_sapiens.GRCh38.cds.all.fa release-110) from Ensembl and the predicted cDNA sequences from the genome assembly for *X. birchmanni*. Since all teleost fish have undergone an ancient whole genome duplication, there were a substantial number of genes that returned multiple hits from the blast_rec command. We assigned an ortholog as a reciprocal best blast hit if there was a single ortholog, or if there was an ortholog with a lower e-value than other hits. We note that the total number of 1:1 orthogs that we are able to identify is likely impacted by the teleost whole genome duplication.

We downloaded the HuMAP2_IDs of genes that are involved in protein complexes in humans from http://humap2.proteincomplexes.org. We converted these to ensembl id using the MANE database (https://www.ncbi.nlm.nih.gov/refseq/MANE/). The HuMAP2_IDs were then matched to *X. birchmanni* orthologs of these genes and their coordinates in the *X. birchmanni* reference genome. This resulted in a dataset of 5,118 *X. birchmanni* genes where the human ortholog is involved in a protein complex.

With these coordinates in hand, we next evaluated average ancestry in regions containing genes involved in protein complexes compared to expectations by chance. We compared average ancestry of protein complex genes in Chapulhuacanito and Santa Cruz to null datasets. To generate null datasets, we randomly selected 5,118 genes not annotated as being involved in protein complexes, calculated average ancestry, and repeated this process 1,000 times. We repeated this analysis using any gene with a 1:1 ortholog between *X. birchmanni* and humans as the focal set, and again compared ancestry in these regions to 1,000 null datasets, each with 5,118 genes (sampled out of a total of 11,289 genes with 1:1 orthologs).

While we found that genes involved in protein complexes had lower average minor parent ancestry in *X. birchmanni* x *X. cortezi* hybrid populations than average minor parent ancestry across all protein-coding regions, the depletion of minor parent ancestry was not significantly lower than observed for other genes with 1:1 orthologs (Fig. S17). This suggests that the signal we detect is simply a consequence of such genes being under greater evolutionary constraint. However, this does not rule out the possibility that certain protein complexes are especially depleted in minor parent ancestry [20].

*Supporting Information 11. Estimating the length of X. birchmanni chromosomes in centimorgans*

For several of the analyses in the paper, it is useful to consider the length of each chromosome in centimorgans (cM). To generate these estimates, we took advantage of artificial crosses between *X. birchmanni* and *X. malinche* generated in the lab between $F_1$ parents, which yielded a large dataset of 1253 second generation hybrids. Because we expect recombination maps to be conserved across *Xiphophorus* species (see Supporting Information 3), crossover maps give us a more direct (albeit coarser) estimate of the recombination rate, that can be used to convert physical to genetic distance.

Because $F_1$ individuals are heterozygous for ancestry at every location in their genome, any ancestry transitions observed in $F_2$ hybrids (apart from those attributable to error), reflect crossover events that occurred during meiosis in the $F_1$ parents. We were thus interested in identifying the number of ancestry transition events per individual on each chromosome to estimate the genetic length of the chromosomes. We also wanted to exclude switch errors where possible, which will have the effect of inflating our estimates of genetic length. We identified and excluded ancestry switches that occurred within 250 kb of each other, as we expect only one crossover per chromosome per meiosis in $F_2$ hybrids and such excess crossovers likely represent errors. We excluded individuals with an extremely high numbers of crossovers across the genome (>75; 12 individuals).

Following this filtering, we recorded the number of crossovers (i.e. ancestry transitions) per individual per chromosome. For each chromosome, we calculated the average number of crossovers between the first and last basepair on the assembled chromosome. Since each $F_2$ individual is the product of two meiosis events, we divided this number by two. We treated this value as the average frequency of recombination events across the length of this chromosome during meiosis. By multiplying this number by 100, we arrived at an estimate of the probability of recombination over the chromosome per meiosis, or the length of each chromosome in centimorgans. We used these estimates of the length of the chromosome in centimorgans, combined with our linkage-disequilibrium maps to roughly convert estimates of $\rho$/bp to genetic distance, and determine intervals of a given genetic distance across the 24 chromosome (e.g 0.1 cM windows).

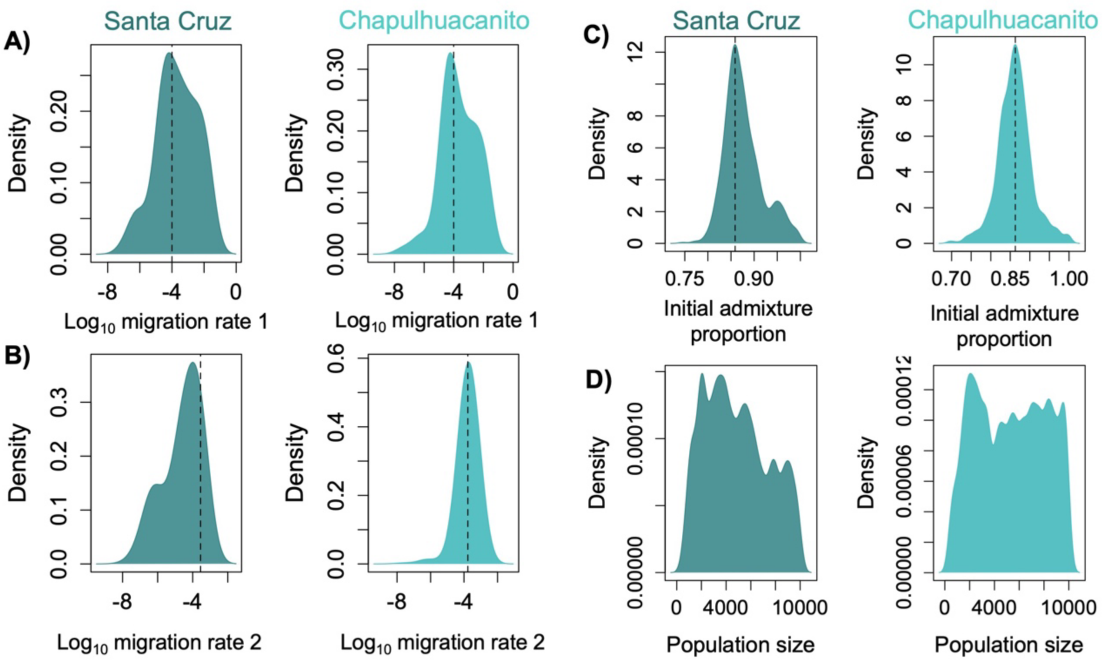


**Fig. S1.** Posterior distributions for all demographic parameters from ABCreg analysis (except generations since initial hybridization; see Fig. 1) for Santa Cruz and Chapulhuacanito populations. We obtain well-resolved posterior distributions for most parameters, including migration rate from *X. cortezi* parent (**A**), migration rate from the *X. birchmanni* parent (**B**), and initial admixture proportion (**C**), but not for hybrid population size (**D**) where we essentially recover the prior distribution (a uniform distribution from 2-10,000). For demographic parameters with well-resolved posterior distributions, the dotted lines indicated the maximum a posteriori estimate for that parameter.

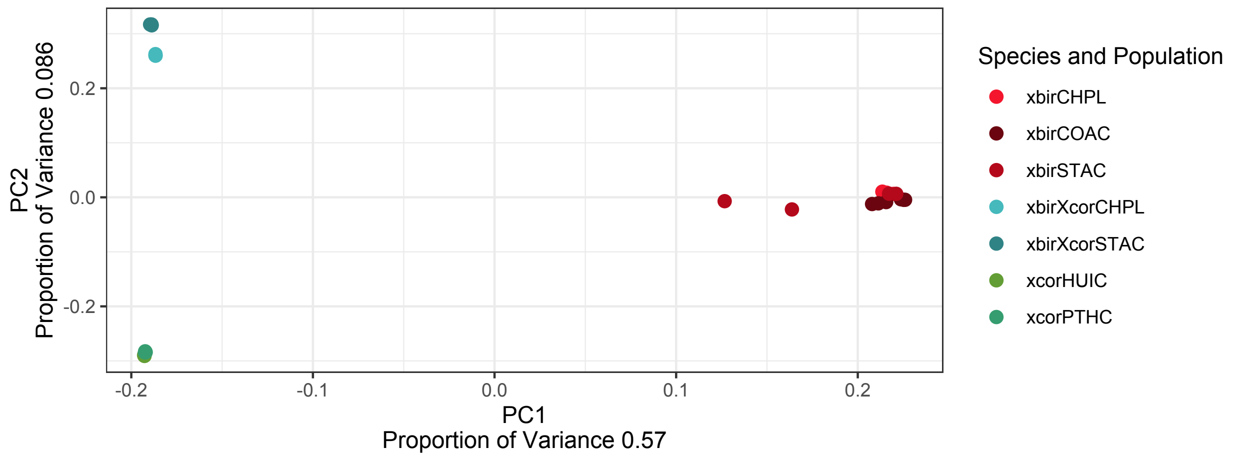


**Fig. S2.** Results of principal component analysis of individuals sequenced to high coverage from Santa Cruz and Chapulhuacanito as well as from allopatric parental populations including all variable sites across the genome. Analyses in the main text show PCAs based on particular ancestry tracts (i.e. homozygous *X. cortezi* or homozygous *X. birchmanni*; Fig. 2). This analysis includes all variant sites from all ancestry tracts. *X. cortezi* and *X. birchmanni* separate along PC1. Hybrids cluster close to *X. cortezi* along PC1, as expected given that they derive >70% of their genome from *X. cortezi*. We include pure *X. birchmanni* individuals found sympatrically with hybrids from Santa Cruz and Chapulhuacanito (xbirCHPL and xbirSTAC), as well as parental individuals from allopatric populations (xbirCOAC, xcorHUIC and xcorPTHC). Hybrids from the Santa Cruz and Chapulhuacanito populations separate from each other in this analysis (xbirXcorSTAC and xbirXcorCHPL codes in the plot respectively).

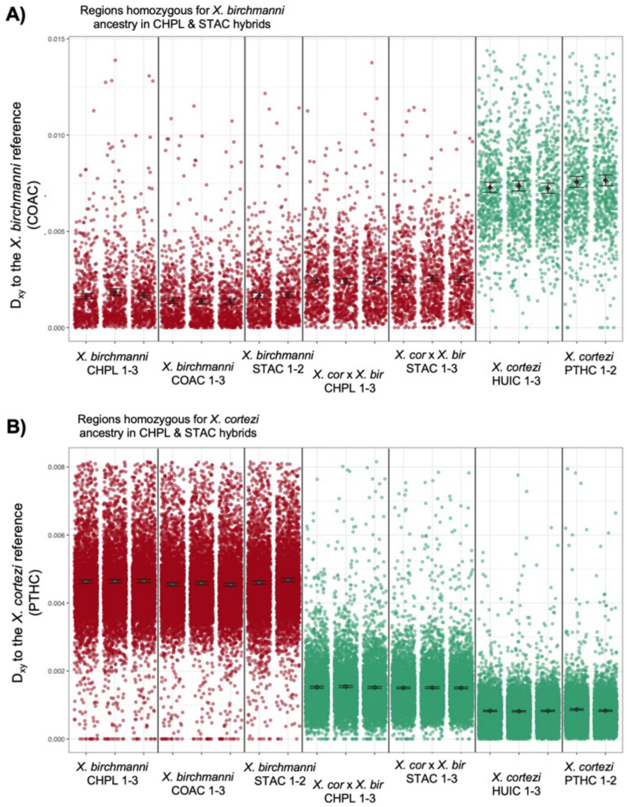


**Fig. S3.** Additional analyses of population genetic patterns in high coverage whole genome sequence data from Santa Cruz and Chapulhuacanito. **A**) We identified homozygous *X. birchmanni* ancestry tracts found in both Santa Cruz (STAC) and Chapulhuacanito (CHPL). We also subsampled these same regions from sympatric and allopatric pure parental individuals (*X. birchmanni* at COAC, CHPL, and STAC, *X. cortezi* at HUIC and PTHC). We found that Dxy to the reference *X. birchmanni* sequence was similar across the sequences analyzed for *X. birchmani* tracts from hybrids and for those tracts from pure *X. birchmanni* individuals. As expected, *X. cortezi* populations had elevated Dxy to the *X. birchmanni* reference. **B**) Similar patterns are observed for *X. cortezi* ancestry tracts analyzed with the same approach. *X. birchmanni*-derived regions had uniformly elevated Dxy compared to the *X. cortezi* reference (derived from the Puente de Huichihuyan or PTHC populations). Notably, homozygous *X. cortezi* ancestry tracts in hybrid individuals (*X. cor* x *X. bir* STAC and CHPL) have slightly but significantly higher Dxy to the *X. cortezi* reference sequence hinting that the source population in both hybrid populations was somewhat diverged from the allopatric *X. cortezi* populations we sampled (consistent with expectations from geography). For both panels, colored points show the raw data and the black point and whiskers shows the mean ± two standard errors of the mean.

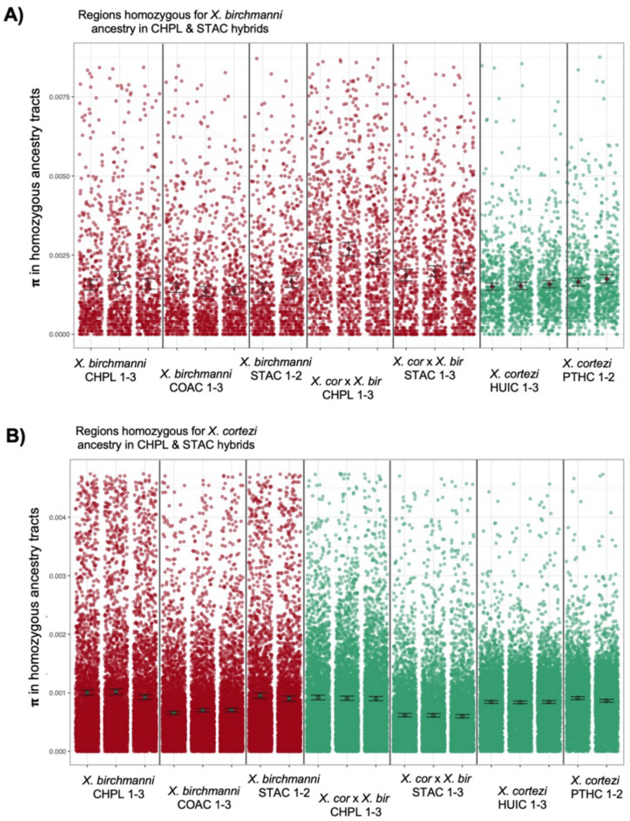


**Fig. S4.** Additional analyses of population genetic patterns in high coverage whole genome sequence data from Santa Cruz and Chapulhuacanito focusing on patterns of nucleotide diversity. In **A)**, we identified regions of the genome that were homozygous for *X. birchmanni* ancestry in the three deep-sequenced hybrids from Chapulhuacanito and the three from Santa Cruz. We subset these regions from the hybrids as well as allopatric parental populations and sympatric *X. birchmanni* individuals (*X. birchmanni* at COAC, CHPL, and STAC, *X. cortezi* at HUIC and PTHC). We find that pure *X. birchmanni* individuals have similar estimated π in these ancestry tracts, whereas Chapulhuacanito and to a lesser extent Santa Cruz have elevated π. In **B)**, we identified regions of the genome that were homozygous for *X. cortezi* ancestry in the three deep-sequenced hybrids from Chapulhuacanito and the three from Santa Cruz. We subset these regions from the hybrids as well as allopatric parental populations and sympatric *X. birchmanni* individuals. We found that although hybrids at Chapulhuacanito have similar estimated π to the *X. cortezi* parent in these ancestry tracts, hybrids at Santa Cruz have markedly lower estimated π. This may point to differences in the demographic history of Santa Cruz hybrids since the population formed, or in the *X. cortezi* parental population that contributed to the hybrids. For both panels, colored points show the raw data and the black point and whiskers shows the mean ± two standard errors of the mean.

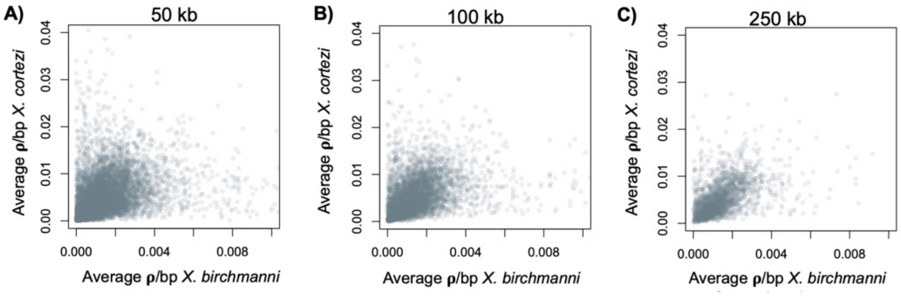


**Fig. S5.** Comparisons of inferred local recombination rates in *X. birchmanni* and *X. cortezi*. Since *Xiphophorus* species lack a PRDM9 ortholog that is active in meiosis and recombine primarily at functional elements that are largely conserved between species (Fig. S6), we expected their recombination maps to be conserved as well. Comparison of local recombination rates across a range of window sizes (**A-C**) confirms this prediction (**A** – Spearman's $\rho = 0.55$; **B** – Spearman's $\rho = 0.57$; **C** – Spearman's $\rho = 0.62$). Given that previous simulations of the *X. birchmanni* recombination map indicated that the expected correlations between the true recombination map and the inferred LD-based recombination map were approximately 0.65 in 50 kb windows [2], this suggests that the two maps may be nearly identical. See Supporting Information 3 for more detail.

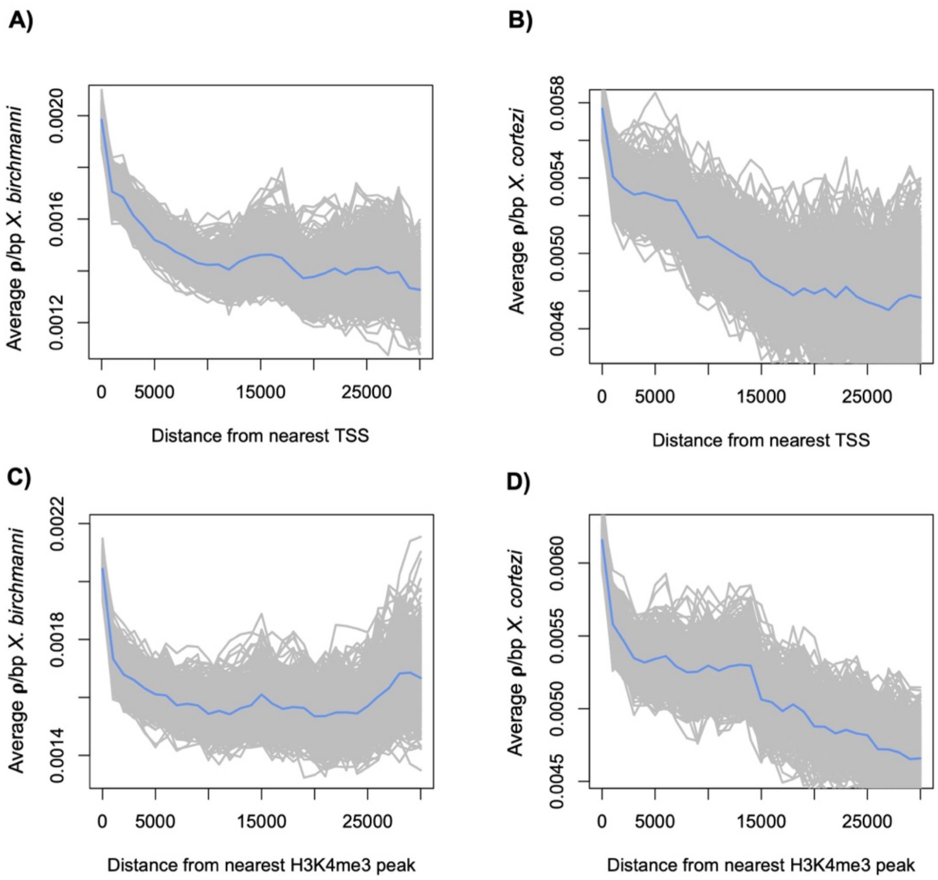


**Fig. S6.** Average recombination rate (ρ/bp) in sliding 5 kb windows as a function of distance of that window from the nearest transcriptional start site (TSS) or H3K4me3 peak in testis tissue in basepairs in *X. birchmanni* (**A**, **C**) and *X. cortezi* (**B**, **D**). In both species, average recombination rates peak near the TSS (**A**, **B**), as frequently observed in species that lack a PRDM9 ortholog active in specifying recombination hotspots. This pattern is thought to be driven by recombination machinery defaulting to the locations of existing H3K4me3 marks in the absence of PRDM9-driven H3K4me3 marks. Indeed, we see that both species have elevated recombination near shared H3K4me3 identified in *X. birchmanni* testis samples (data reanalyzed from [3]; **C**, **D**). Gray lines show individual replicates bootstrap resampling the data, with a total of 500 replicates plotted. Blue line shows the average across simulations.

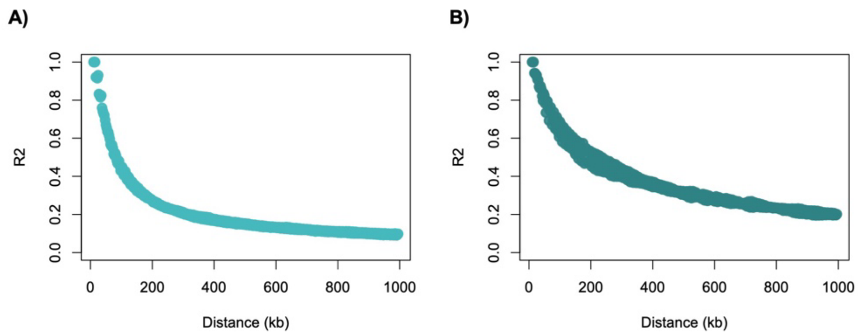


**Fig. S7.** Average decay of admixture linkage disequilibrium among hybrid individuals from the Chapulhuacanito (left) and Santa Cruz (right) hybrid populations. The y-axis shows the average $R^2$ value in sliding 5 kb windows and the x-axis shows the physical distance between ancestry informative sites in the focal window. Both populations asymptote to background levels of admixture LD by approximately 500 kb, although we thin our data by 1 Mb to be conservative (see Methods). We note also that there is an excess of admixture linkage disequilibrium in both populations where the minimum average $R^2$ greatly exceeds one over the number of sampled individuals, particularly in the Santa Cruz population. This is suggestive of population structure, a recent pulse of migration, or assortative mating in these populations (see [8] for data on assortative mating in Santa Cruz).

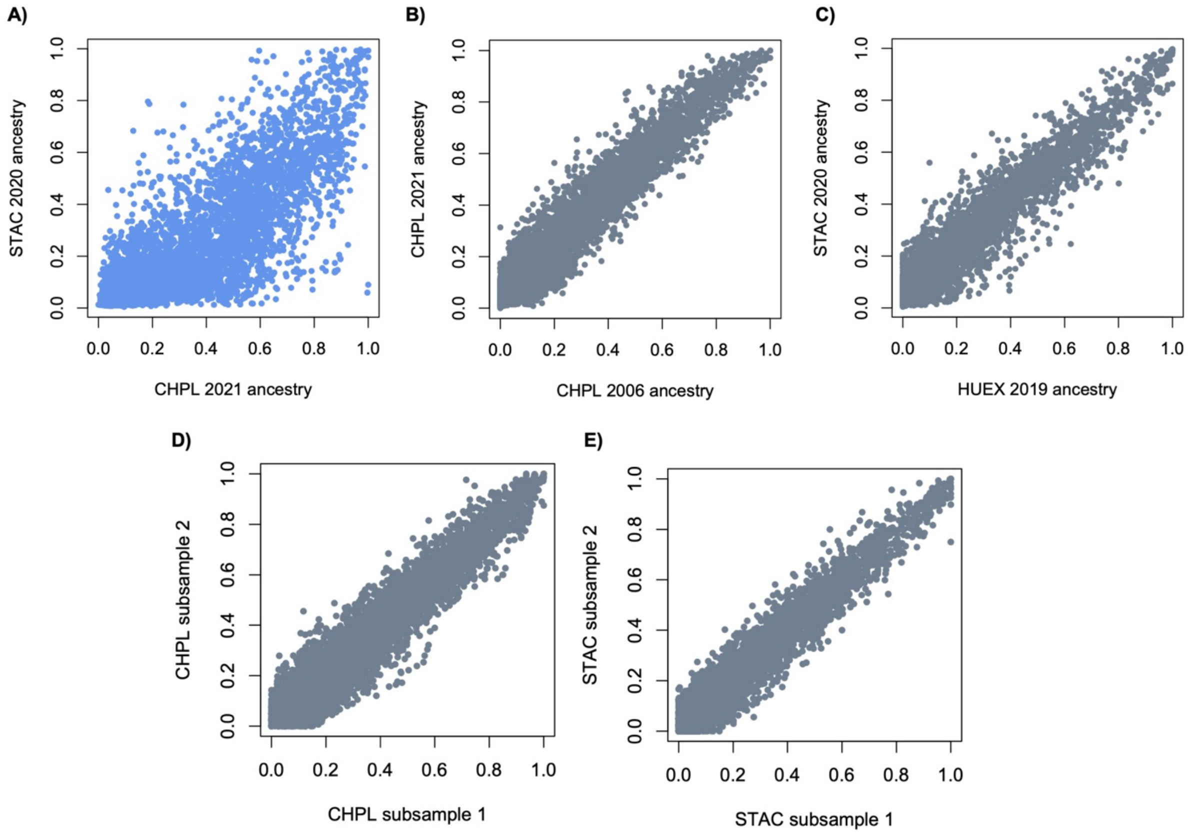


**Fig. S8.** Visualization of cross-population versus within population (or river) correlations in minor parent ancestry. Average minor parent ancestry for matched regions of the genome is plotted on the x and y axes in 100 kb windows. **A**) Average ancestry in Chapulhuacanito (CHPL) population samples from 2021 are plotted against average ancestry in the same windows from the Santa Cruz (STAC) population sampled in 2020. Correlations in local ancestry are high, but windows with high minor parent ancestry in Chapulhuacanito and low minor parent ancestry in Santa Cruz (and vice versa) can be visualized in the plot. By contrast, local ancestry comparisons from the same population but different years (**B**) or two nearby populations in the same river drainage (**C**) are remarkably concordant. **D** and **E** show results from subsampling twenty individuals from the same population and sampling year. Again, correlations between individuals sampled from the same population (**D** – Chapulhuacanito, **E** – Santa Cruz), greatly exceed observed correlations between drainages (**A**).

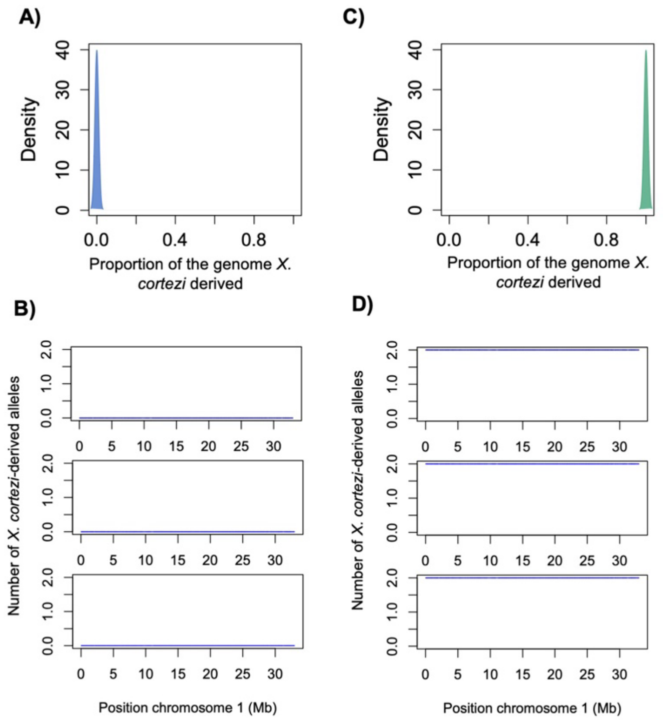


**Fig. S9.** **A)** Analyses of genome-wide ancestry in allopatric *X. birchmanni* individuals demonstrates that *ancestryinfer* correctly infers that these individuals are homozygous for *X. birchmanni* ancestry across the genome. **B)** This result can also be observed by examining local ancestry in the same individuals (three representative individuals plotted here), where entire chromosomes are inferred to have two *X. birchmanni*-derived and no *X. cortezi*-derived haplotypes. **C**) Similarly, ancestry inference in allopatric *X. cortezi* individuals is accurate, with all individuals inferred to be homozygous *X. cortezi* throughout the genome. **D**) This result can also be observed by examining local ancestry in the same individuals (three representative individuals plotted here), where entire chromosomes are inferred to have two *X. cortezi* derived and no *X. birchmanni*-derived haplotypes.

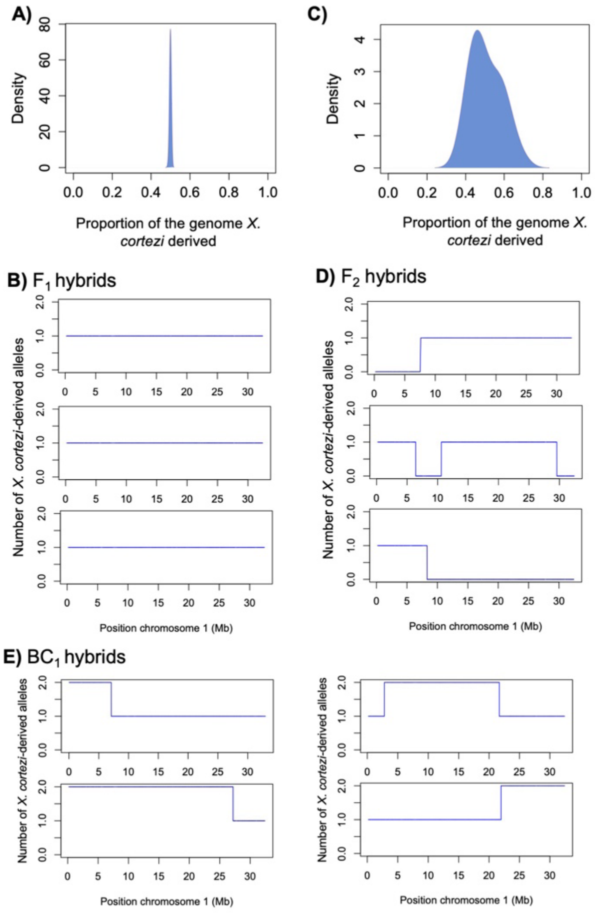


**Fig. S10.** Analyses of genome-wide ancestry in hybrid offspring of known crosses. **A**) $F_1$ hybrids produced in lab are inferred by *ancestryinfer* to have precisely 50% of their genome derived from *X. birchmanni* and 50% from *X. cortezi*, with essentially no variation in ancestry observed, as expected for this cross. **B**) Three representative individuals showing local ancestry across chromosome 1 in $F_1$ hybrids. $F_1$ hybrids are inferred to have one *X. birchmanni* and one *X. cortezi* haplotype across the chromosome, as expected from the cross design. **C**) $F_2$ hybrids produced in lab are inferred to have on average 50% of their genome derived from each parental species, but with substantial variation in genome-wide ancestry induced by recombination between the *X. birchmanni* and *X. cortezi* haplotypes in their $F_1$ parents, followed by independent assortment. **D**) Local ancestry across chromosome 1 for three representative $F_2$ individuals. **E**) Local ancestry across chromosome 1 for four representative $BC_1$ individuals ($F_1$ hybrids crossed to *X. cortezi*). We produced fewer backcross individuals than other cross types so do not plot genome-wide ancestry for these individuals.

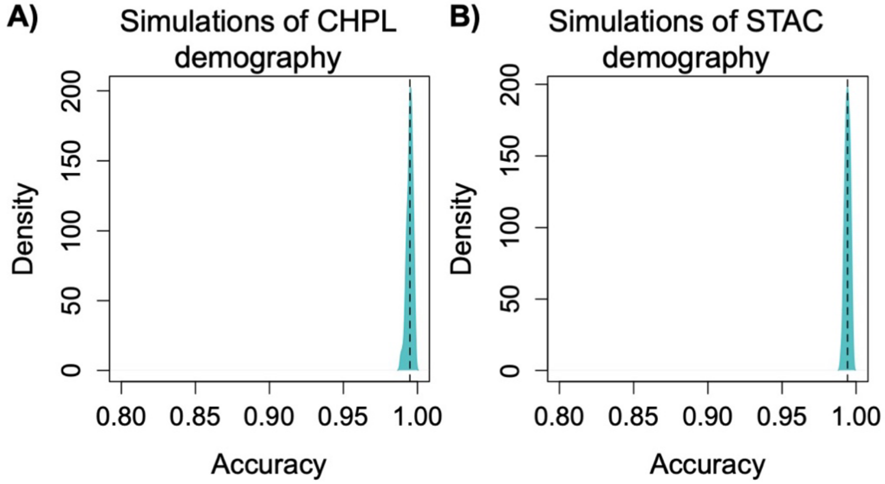


**Fig. S11.** Estimates of error rate using *mixnmatch* simulations. As a complementary analysis to ancestry inference in known crosses, we used the program *mixnmatch* to simulate late generation hybrids with parameters matching the Chapulhuacanito (**A**) and Santa Cruz (**B**) populations to estimate our error rate in the case of late generation hybrids between *X. cortezi* and *X. birchmanni*. **A)** Simulations of Chapulhuacanito hybrids using the MAP estimates of demographic parameters from the posterior distributions generated by ABCreg and with a simulated hybrid population size of 5,000. Accuracy in local ancestry inference for 50 simulated hybrid individuals is shown here. The average error rate in these simulations was 0.3%. **B**) Simulations of Santa Cruz using the MAP estimates of demographic parameters from the posterior distributions generated by ABCreg and with a simulated hybrid population size of 5,000. Accuracy in local ancestry inference for 50 simulated hybrid individuals is shown here. The average error rate in these simulations was 0.4%. As expected, the error rate is slightly higher for simulations of Santa Cruz, since this population is likely to be older and smaller ancestry tracts are more difficult to accurately infer.

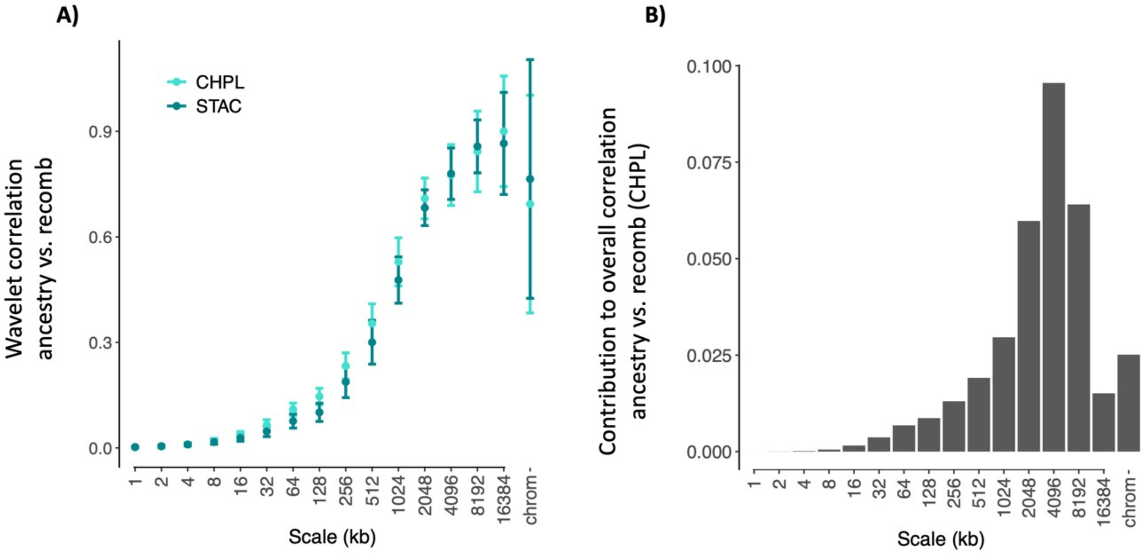


**Fig. S12. A)** Spatial wavelet decomposition of the overall Pearson correlation between inferred minor parent ancestry in Chapulhuacanito (CHPL) and Santa Cruz (STAC) versus the inferred recombination rate, measured at a resolution of 1 kb. **B)** The contribution of a given spatial scale to the overall correlation is a weighted correlation of wavelet coefficients for the two signals at that scale, weighted by the variances in each signal at that scale, also obtained from the discrete wavelet transform. We show this decomposition for Chapulhuacanito and note that the general pattern is highly similar for Santa Cruz (STAC). The contributions sum to the total Pearson correlation calculated at the finest resolution of measurement (1 kb). For simplicity we omit 'scaling' variances which are leftover variance due irregularity of chromosome lengths (for further discussion see [23]).

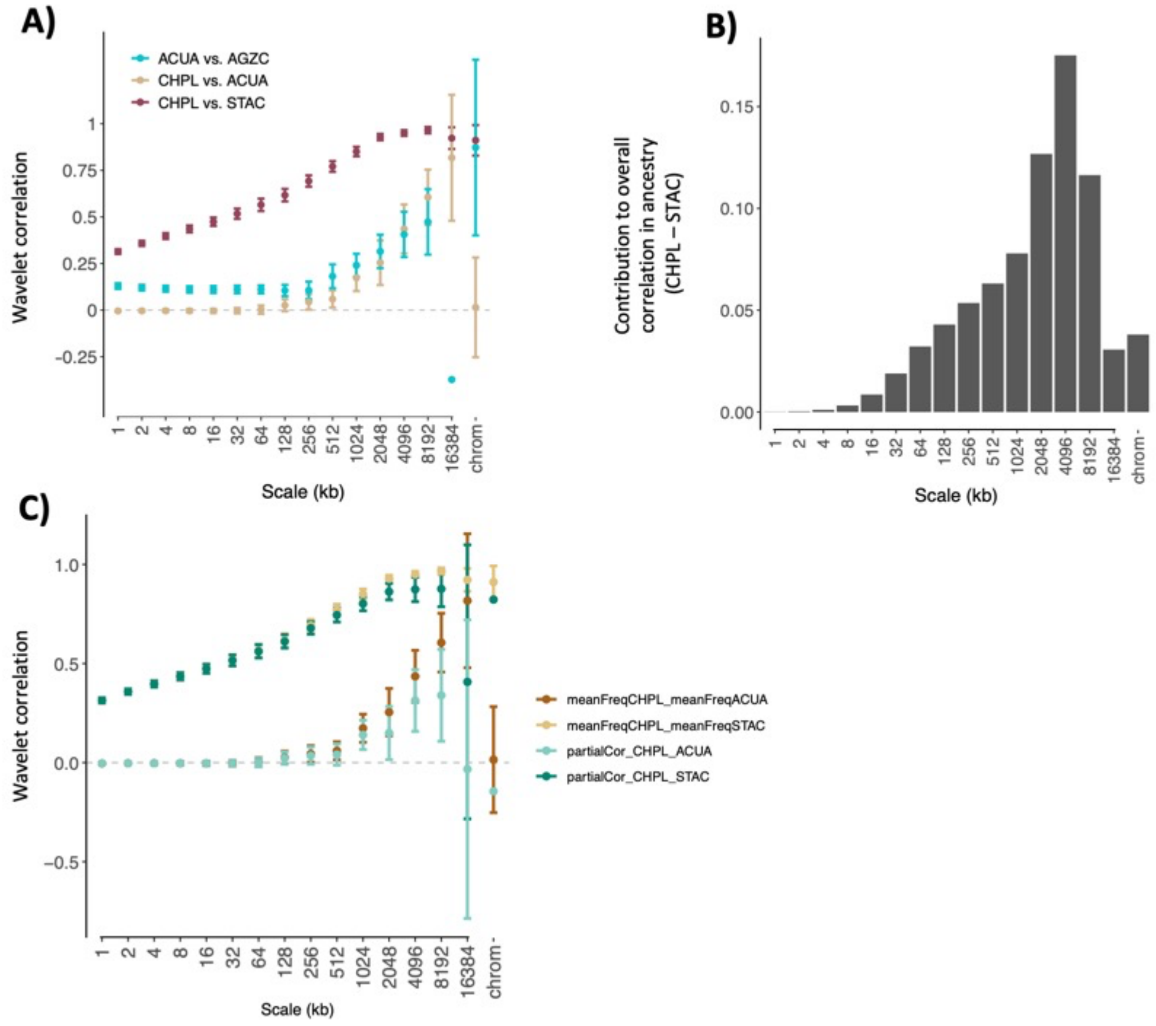


**Fig. S13. A)** Wavelet correlations between minor parent ancestry proportion in cross population comparisons between hybrids derived from the same hybridizing pair of *X. birchmanni* x *X. cortezi* (CHPL vs. STAC), the same hybridizing pair of *X. birchmanni* x *X. malinche* populations (ACUA vs AGZC), and between hybrids from different hybridizing pairs (CHPL vs. ACUA). Points are weighted averages across chromosomes with error bars representing 95% jackknife confidence intervals. For visualization, we omit the confidence interval for the wavelet correlation of ancestry in ACUA vs. AGZC at the largest scale, since it is large and overlaps zero. See discussion in Supporting Information 7 regarding fine scale correlations (e.g. 1 kb). **B)** Here, the correlations in **A** are weighted by the variances in ancestry in each population at a given scale to reflect the contribution of that scale to the overall correlation. **C)** Partial wavelet correlations of minor parent ancestry between populations after accounting for variation in recombination. At each scale, we compute a wavelet correlation of the residuals for each population from a linear model with recombination as a predictor. Using these values, we find a comparatively larger reduction in comparisons across hybrid population types (i.e. CHPL vs.

ACUA). This would be consistent with a larger portion of the ancestry correlation in these comparisons being driven by the shared effects of recombination.

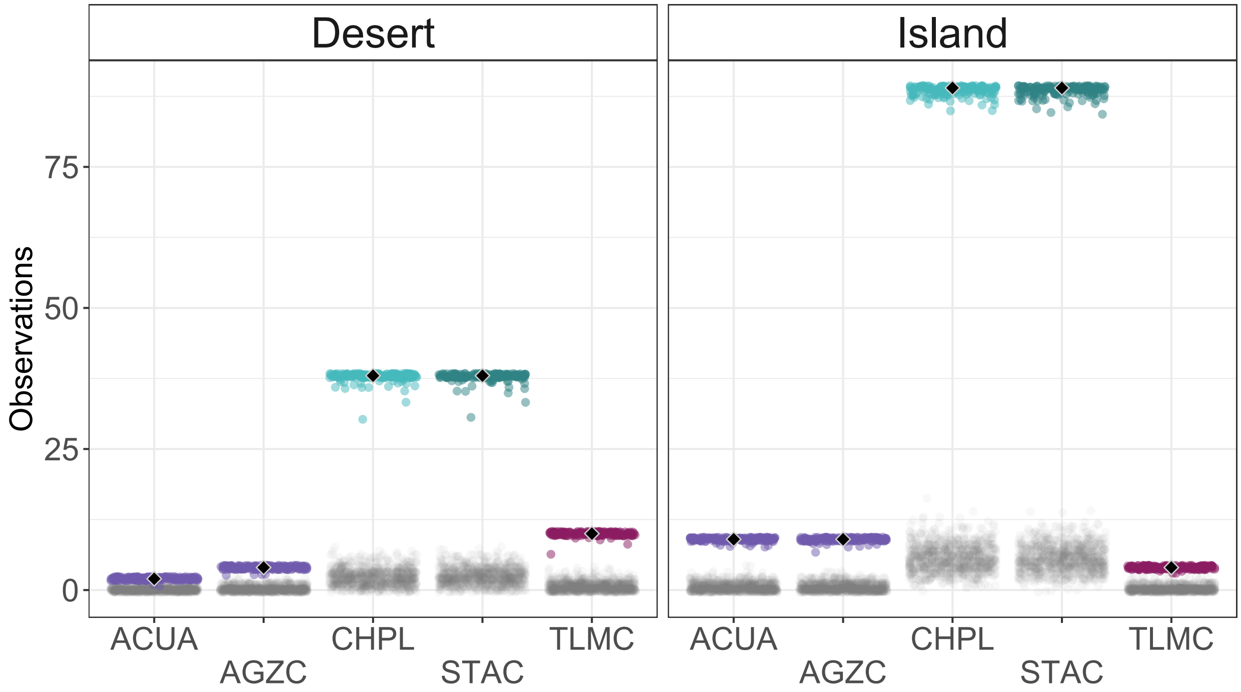


**Fig. S14.** Observed shared minor parent deserts and islands across hybrid population types. Minor parent deserts and islands were identified in the *X. birchmanni* x *X. cortezi* hybrid populations Chapulhuacanito (CHPL) and Santa Cruz (STAC). We then asked whether these regions overlapped with minor parent deserts or islands found in *X. birchmanni* x *X. malinche* hybrid populations (Acuapa – ACUA, Aguazarca – AGZC, and Tlatemaco – TLMC). Black diamonds indicate the observed number of overlaps between minor parent deserts and islands in any *X. birchmanni* x *X. malinche* population and the *X. birchmanni* x *X. cortezi* hybrid populations (with the number shared between Chapulhuacanito and Santa Cruz shown in blue for comparison). Colored points show the number of shared deserts or islands from replicates jack-knife bootstrapping the genome in 10 cM windows. Gray points show null expectations for each comparison (see Methods). As expected, we observe many more shared deserts and islands among *X. birchmanni* x *X. cortezi* hybrid populations than between *X. birchmanni* x *X. cortezi* and *X. birchmanni* x *X. malinche* hybrid populations.

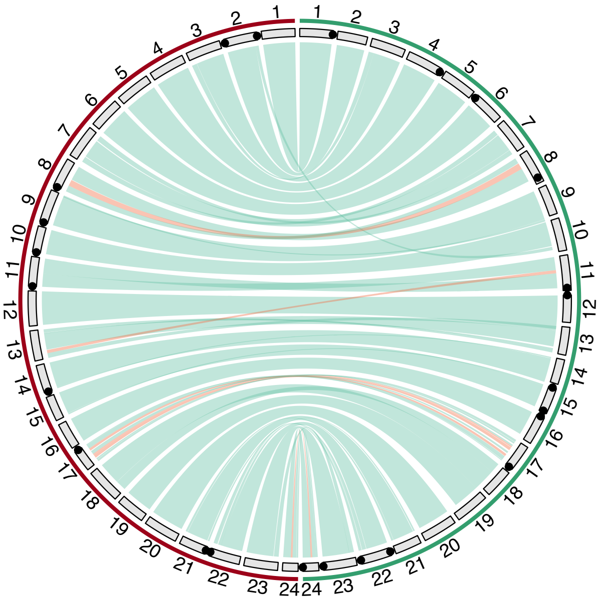


**Fig. S15.** Circos plot of the new *X. cortezi* genome assembly (green) relative to *X. birchmanni* (red). Inversions between the two species are highlighted by light red lines connecting chromosomes, while co-linear regions are connected by light green lines (based on minimap2 alignments). Gray boxes show schematics of each chromosome. Black circles on the chromosome schematic indicate locations on the chromosomes where telomeric sequences were detected (using seqtk telo). The Circos plot was generated by the R package circulize. Note that several putative translocations are identified, which we treat with caution in the absence of Hi-C data for *X. cortezi*.

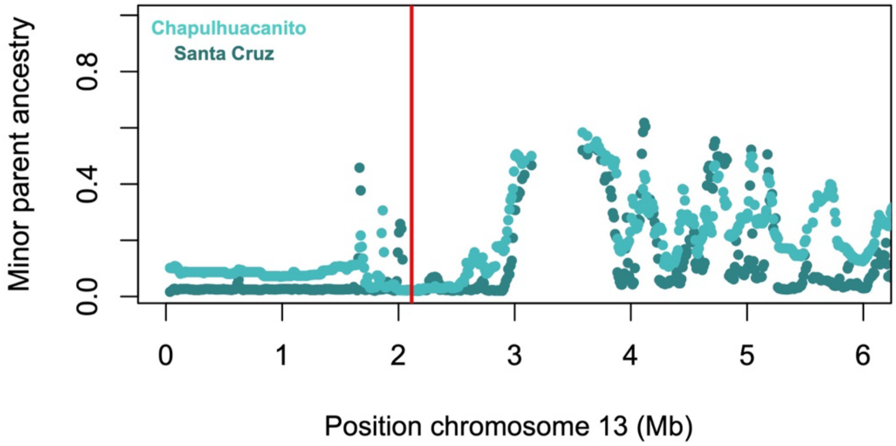


**Fig. S16.** Plot of average minor parent (*X. birchmanni*) ancestry across the first six Mb of chromosome 13 in both Santa Cruz and Chapulhuacanito with the location of *ndufs5* highlighted by the red solid line. *ndufs5* is involved in a known mitonuclear incompatibility (see main text) but the region surrounding it was not identified as a shared desert (found as a desert in Chapulhuacanito but not Santa Cruz). However, examination of local ancestry indicates that *X. birchmanni* ancestry at *ndufs5* (red line) is rare in both populations, but was slightly higher than the lowest 5% quantile for Santa Cruz (Lowest 5% quantile for *X. birchmanni* ancestry is 2.2% in Santa Cruz, versus 2.3% *X. birchmanni* ancestry observed at *ndufs5*).

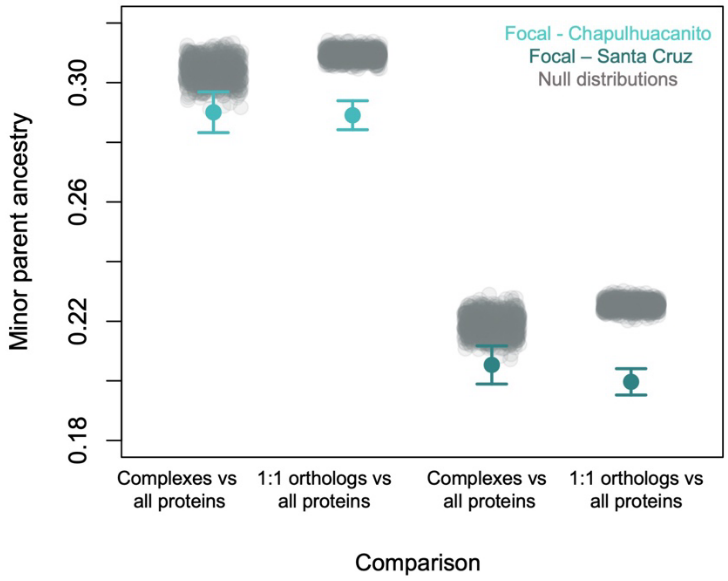


**Fig. S17.** Comparisons of minor parent ancestry across genes of different annotations in the two hybrid populations. In the "Complexes vs all proteins" comparison, we calculated average minor parent ancestry in genes involved in protein complexes (see Supporting Information 10) in both Chapulhuacanito and Santa Cruz (colored points and whiskers), compared to null datasets generated from randomly sampling matched sets of protein coding genes (gray points). Observed minor parent ancestry in these proteins is slightly but significantly lower than the null distribution. We next considered the possibility that this signal was driven by genes involved in protein complexes being highly conserved. We identified all genes that had a single reciprocal best blast hit with human proteins ("1:1 orthologs"), excluded genes involved in protein complexes, and repeated the analysis (see Supporting Information 10). We found that these genes were similarly depleted in minor parent ancestry relative to the null, suggesting that the signal we observe with genes in protein complexes is driven by conservation, not directly by their role in protein-protein interactions.

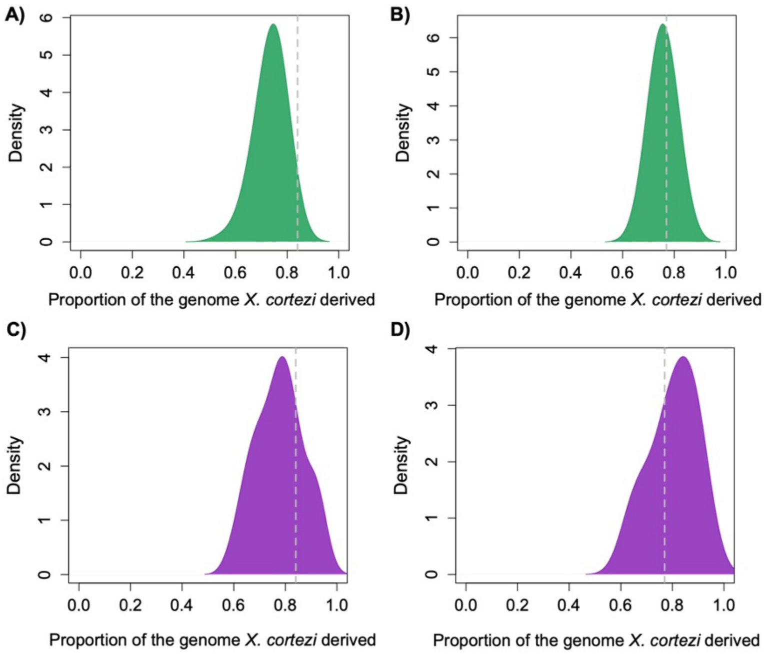


**Fig. S18.** Results of admix'em simulations with selection. Plots show genome-wide admixture proportions at the end of simulations described in Supporting Information 4. **A** and **C** show simulations using demographic parameters inferred for the Santa Cruz population from ABCreg, with selection on 40 recessive hybrid incompatibilities (**A**) or on 40 hybrid incompatibilities with a range of dominance coefficients (**C**). The dotted gray line shows the observed admixture proportion in Santa Cruz. **B** and **D** show simulations using demographic parameters inferred for the Chapulhuacanito hybrid population from ABCreg, with selection on 40 recessive hybrid incompatibilities (**B**) or on 40 hybrid incompatibilities with a range of dominance coefficients (**D**). The dotted gray line shows the observed admixture proportion in Chapulhuacanito. For all sets of admix'em simulations, we had to modify the initial admixture proportion from the posterior distributions inferred by ABCreg because simulation selection dramatically shifts the initial admixture proportion (see Supporting Information 4). As a result, we wanted to confirm that after simulations of selection with admix'em, the final admixture proportions roughly matched those observed in the Santa Cruz and Chapulhuacanito hybrid populations.

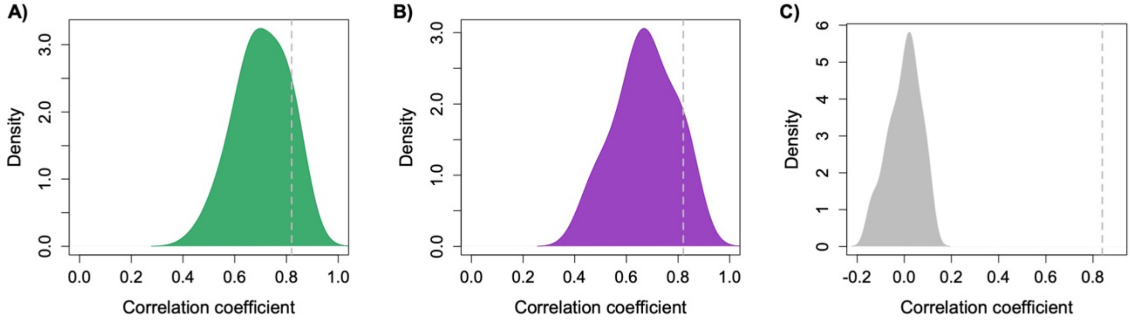


**Fig. S19.** Results of admix'em simulations modeling shared selection in two independently formed hybrid populations. For each simulation, we drew demographic parameters from the posterior distributions inferred by ABCreg for Santa Cruz and Chapulhuacanito. We simulated two hybrid populations using these demographic parameters as described in Supporting Information 4. In one set of simulations, we implemented selection on 40 pairs of recessive hybrid incompatibilities, with selection coefficients drawn from an exponential distribution with a mean of 0.6. The observed cross-population correlations in ancestry in 250 kb windows for 50 replicates of this simulation scenario are shown in (**A**). In another set of simulations, we implemented selection on 40 pairs of hybrid incompatibilities as before, except that we drew dominance coefficients from a uniform distribution of 0-0.5 and selection coefficients from an exponential distribution with a mean of 0.4. The observed cross-population correlations in ancestry in 250 kb windows for 50 replicates of this simulation scenario are shown in (**B**). The inferred cross-population correlation coefficient in 250 kb windows between Santa Cruz and Chapulhuacanito in shown by the dotted gray line. Although the observed value falls in the upper range of the distribution inferred from simulations, these results indicate that selection, combined with the inferred demographic history of these two populations, can drive the high correlations in local ancestry we observe. **C)** Neutral simulations with no selection implemented are plotted for comparison (see Supporting Information 4).

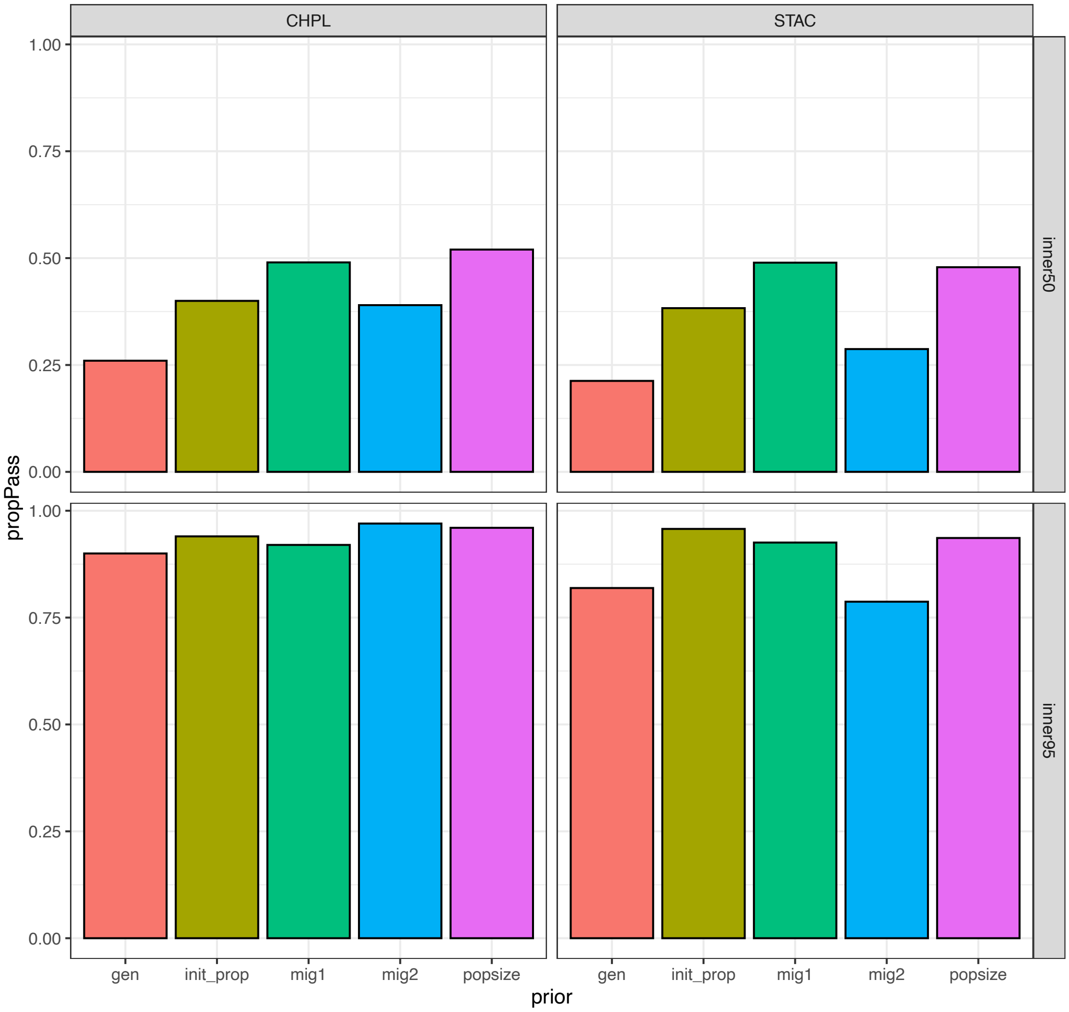


**Fig. S20**. Analysis of expected performance of ABCreg. To evaluate the expected performance of ABCreg, we randomly sampled 100 simulations generated for ABC demographic inference. We treated the summary statistics from each simulation as if it were the real data and ran ABCreg. We asked for a given simulation, whether the true value for the focal parameter fell within the 50% quantile of the posterior distribution generated by ABCreg (top) or the 95% quantile (bottom). Plotted are the proportion of simulations where the true value for each parameter fell within the 50% or 95% quantile of the posterior distribution generated by ABCreg. For all parameters, the true value is very likely to fall in the 95% quantile of the posterior distribution generated by ABCreg.

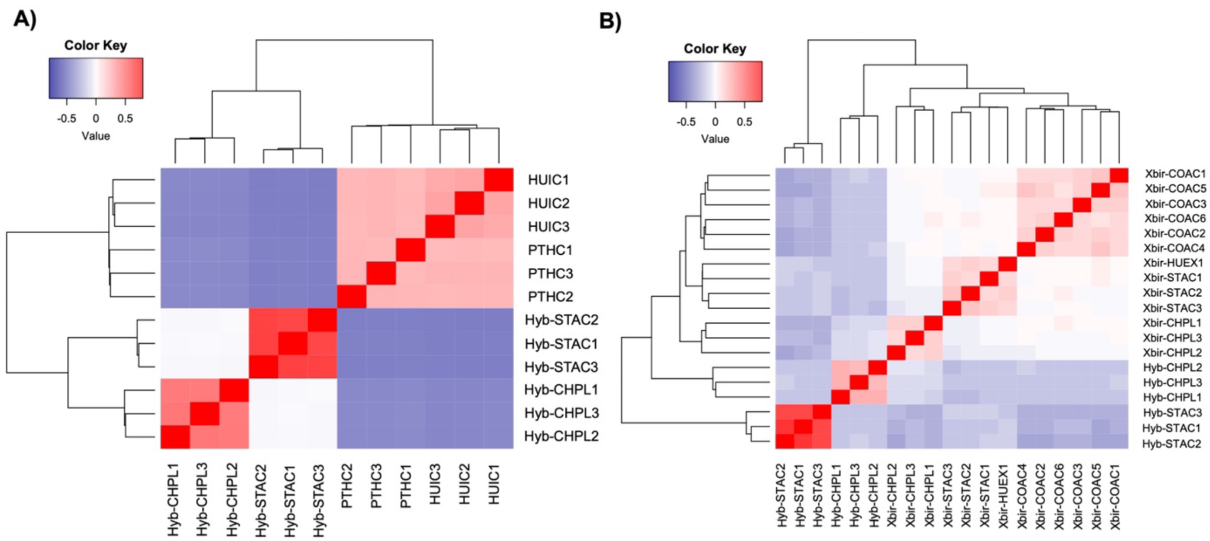


**Fig. S21.** Heatmap of the genetic relatedness matrices across different subsets of samples and regions of the genome. Analysis was performed with GCTA (see Methods). **A)** Analysis of *X. cortezi* ancestry tracts in three hybrids from the Santa Cruz (Hyb-STAC) population and three hybrids from Chapulhuacanito (Hyb-CHPL) population along with the same regions sampled from individuals from the two parental populations, Huichihuyan (HUIC) and Puente de Huichihuyan (PTHC). Cooler colors indicate correlations in the genetic relatedness matrix less than zero and warmer colors indicate positive correlations in the genetic relatedness matrix. Overall, we do not see evidence of genetic relatedness across hybrid populations but substantial evidence of relatedness within populations, with stronger correlations in comparisons between Santa Cruz individuals. High relatedness between pure *X. cortezi* in Huichihuyan and Puente de Huichihuyan is not surprising since these populations occur on the same river. **B)** Results of the same analysis, but for ancestry tracts that were homozygous for *X. birchmanni* in the three high-coverage hybrids from Santa Cruz (Hyb-STAC) and Chapulhuacanito (Hyb-CHPL), as well as the same regions sampled from pure *X. birchmanni* from the Río Santa Cruz (Xbir-STAC, Xbir-HUEX) and Chapulhuacanito (Xbir-CHPL) and from a pure *X. birchmanni* source population (Xbir-COAC). Overall, the results for hybrid populations are concordant with the analysis in **A**, with evidence of relatedness within but not between hybrid populations. However, we do observe more relatedness across *X. birchmanni* individuals from different rivers than we expected *a priori*. See Supporting Information 2 for a discussion of these results.

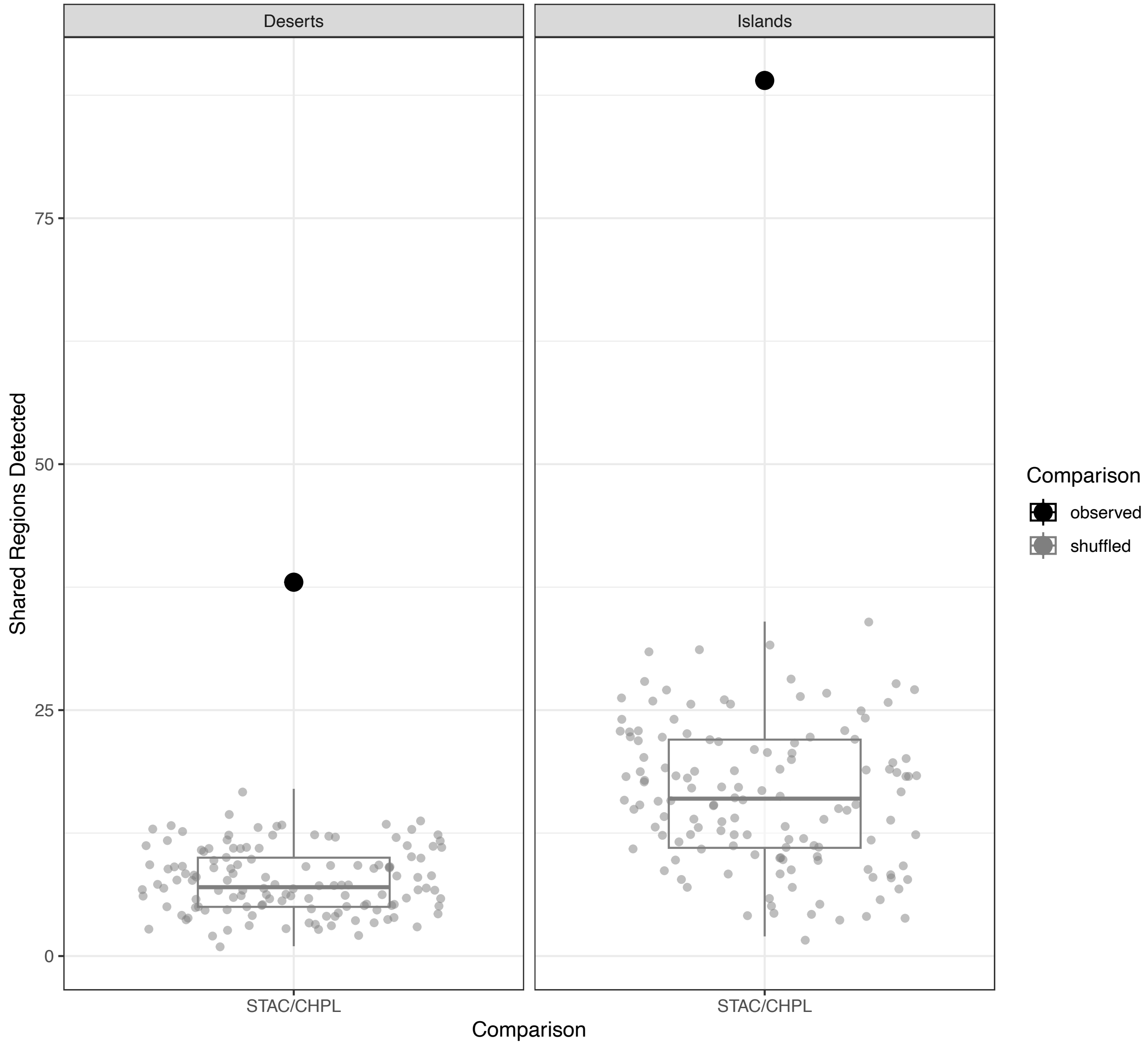


**Fig. S22.** The number of shared deserts (left) and islands (right) between the Santa Cruz (STAC) and Chapulhuacanito (CHPL) populations compared to expectations by chance, using a different permutation approach that preserves the structure of local ancestry variation along the genome (see Methods). Gray points and boxplots indicate expectations from permutations, black points indicate the observed data.

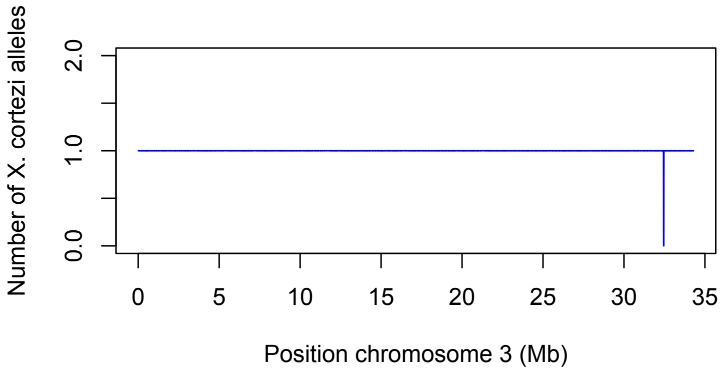


**Fig. S23.** Example of a likely error in local ancestry inference detected in an $F_1$ hybrid between *X. birchmanni* and *X. cortezi* on chromosome 3. In this individual, the local ancestry call switches from heterozygous for ancestry to homozygous *X. birchmanni* for 11 kb. While errors like these are occasionally detected in our datasets of parental and $F_1$ hybrid individuals, we estimate our overall error rates for these individuals to be ~0.1% per ancestry informative site.

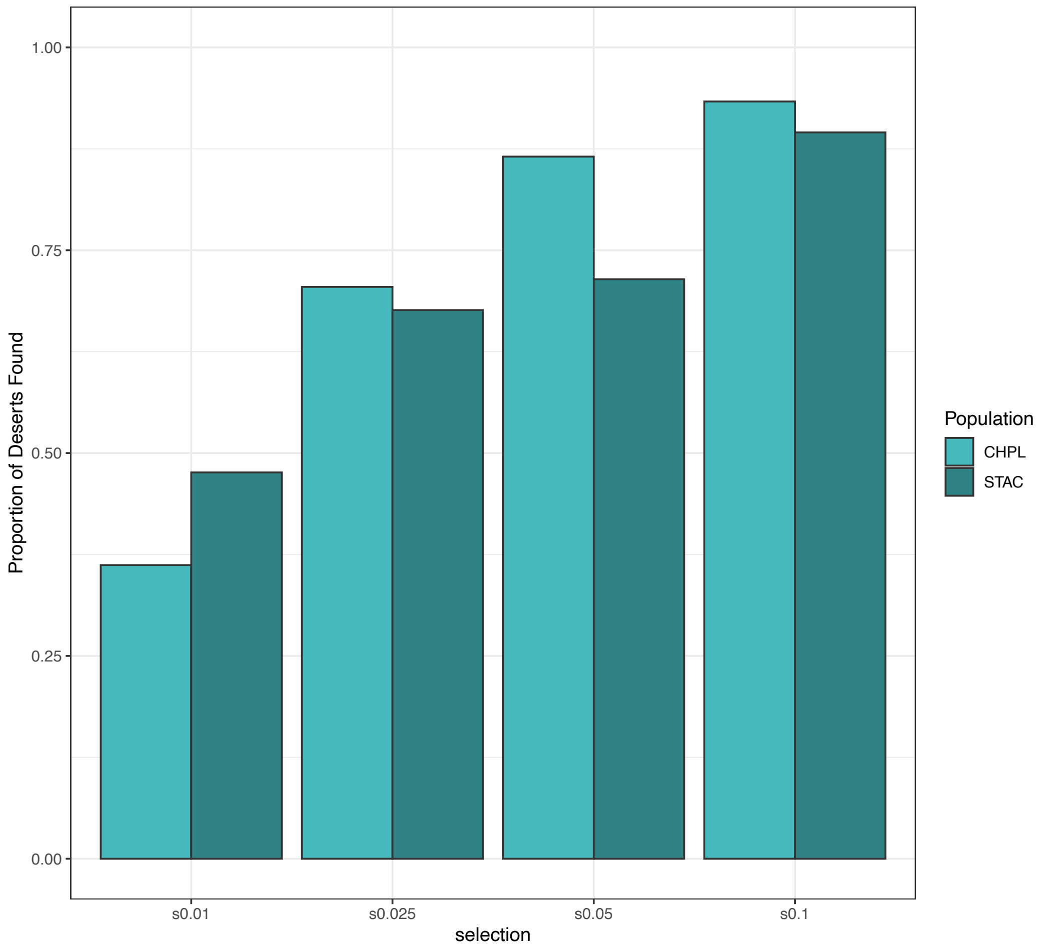


**Fig. S24.** Simulations of power to detect selected sites in populations matching the demographic history of Santa Cruz (STAC) and Chapulhuacanito (CHPL) populations. For each population and selection coefficient (s=0.01-0.1), we performed 100 pairs of simulations of selection using admix'em. For each pair of simulations, we randomly identified the location of the site under selection and then performed simulations with demographic parameters drawn from the ABCreg posterior distributions for Santa Cruz and Chapulhuacanito, respectively. We then identified minor parent deserts as we had for the real data (see Methods), and determined the proportion of time over 100 simulations that we correctly identified the ancestry desert in each population.

**Table S1.** Transposable element classes annotated in *X. birchmanni* and *X. cortezi* with more than a 10% copy number difference in the PacBio HiFi genome assemblies of the two species.

| **Element annotation** | ***X. birchmanni* copy number** | ***X. cortezi* copy number** |
|---|---|---|
| DNA:CMC-EnSpm | 4633756 | 4175251 |
| DNA:Crypton-S | 81900 | 64962 |
| DNA:Dada | 24369 | 21879 |
| DNA:Kolobok-Hydra | 26334 | 22327 |
| DNA:P | 502814 | 440376 |
| DNA:PiggyBac | 542091 | 479037 |
| DNA:RC | 8159880 | 7159251 |
| DNA:Sola-3 | 196508 | 110878 |
| DNA:Zisupton | 351011 | 317145 |
| LINE:I | 2377386 | 2142649 |
| LINE:L1-Tx1 | 1853402 | 1579130 |
| LINE:Proto2 | 268094 | 233548 |
| SINE:tRNA-L1 | 1347298 | 1184614 |
| LTR:ERV-Foamy | 2814375 | 1289681 |
| LTR:ERVK | 388961 | 318244 |
| LTR:ERVL | 9367 | 7272 |
| LTR:Gypsy | 4993068 | 4287995 |
| LINE:RTE-X | 31866 | 27078 |
| DNA:Crypton | 10491 | 8429 |
| DNA:TcMar-Ant1 | 1560 | 1218 |
| DNA:CMC-Transib | 384060 | 439064 |
| DNA:Ginger-2 | 513 | 2570 |
| DNA:Novosib | 16452 | 19915 |
| DNA:PIF-Spy | 3542 | 4456 |
| DNA:TcMar-Mariner | 132570 | 192627 |
| LINE:I-Jockey | 164819 | 258738 |
| LINE:Penelope | 486258 | 601090 |
| LINE:R2-NeSL | 70333 | 77645 |
| LINE:RTE-BovB | 4146214 | 4593954 |
| SINE:tRNA-V-CR1 | 30157 | 34081 |
| LTR:Copia | 644541 | 721754 |
| LINE:Tad1 | 42067 | 46962 |

**Table S2.** Inversions that differentiate *X. birchmanni* and *X. cortezi* based on our PacBio HiFi assemblies. Genome assemblies were aligned with minimap2 and all inversions greater than 100 kb were annotated.

| **Chromosome** | **Start coordinate (*X. birchmanni*)** | **End coordinate (*X. birchmanni*)** |
|---|---|---|
| chr-07 | 6637497 | 7241732 |
| chr-08 | 6542350 | 12967418 |
| chr-08 | 28401987 | 29237938 |
| chr-08 | 13072933 | 13389204 |
| chr-17 | 4580825 | 7858932 |
| chr-17 | 13620980 | 16130397 |
| chr-17 | 7898896 | 9230698 |
| chr-18 | 34136089 | 34325628 |
| chr-24 | 5850012 | 7595198 |

**Table S3.** MAP estimate and 95% percent confidence intervals for posterior distributions generated by ABCreg for each demographic parameter of interest.

| **Parameter (prior)** | **Chapulhuacanito 2.5% quantile** | **Chapulhuacanito MAP estimate** | **Chapulhuacanito 97.5% quantile** | **Santa Cruz 2.5% quantile** | **Santa Cruz MAP estimate** | **Santa Cruz 97.5% quantile** |
|---|---|---|---|---|---|---|
| Population size (2-10,000) | 734 | 3113 | 9760 | 941 | 5652 | 9651 |
| Generations since admixture (10-400) | 104 | 137 | 317 | 218 | 263 | 383 |
| Migration – species 1 (0-3%) | 0 | 0.017% | 2.4% | 0 | 0.028% | 2.3% |
| Migration – species 2 (0-3%) | 0 | 0.004% | 0.11% | 0 | 0.004% | 0.05% |
| Initial admixture proportion (0.5-1) | 0.76 | 0.86 | 0.96 | 0.81 | 0.86 | 0.97 |

**Table S4.** Results of the analysis of the correlation between local minor parent ancestry in windows of varying size (100 – 500 kb reported here) versus the average recombination rate in that window. Since both ancestry and recombination rate are spatially correlated along the genome, for each analysis, we thinned the data to include one window per Mb, such that all analyses presented below include the same number of windows. Population names in italics indicate the focal samples primarily discussed in the main text.

| **Population** | **Window size** | **Spearman's ρ** | **Thinned p-value** |
|---|---|---|---|
| *Santa Cruz 2020* | 100 kb | 0.50 | $<10^{-100}$ |
| *Chapulhuacanito 2021* | 100 kb | 0.55 | $<10^{-56}$ |
| Chapulhuacanito 2003 | 100 kb | 0.50 | $<10^{-43}$ |
| Chapulhuacanito 2006 | 100 kb | 0.51 | $<10^{-47}$ |
| Chapulhuacanito 2017 | 100 kb | 0.55 | $<10^{-55}$ |
| Huextetitla 2003 | 100 kb | 0.50 | $<10^{-44}$ |
| Huextetitla 2019 | 100 kb | 0.49 | $<10^{-43}$ |
| *Santa Cruz 2020* | 250 kb | 0.58 | $<10^{-66}$ |
| *Chapulhuacanito 2021* | 250 kb | 0.58 | $<10^{-65}$ |
| Chapulhuacanito 2003 | 250 kb | 0.57 | $<10^{-61}$ |
| Chapulhuacanito 2006 | 250 kb | 0.59 | $<10^{-66}$ |
| Chapulhuacanito 2017 | 250 kb | 0.60 | $<10^{-70}$ |
| Huextetitla 2003 | 250 kb | 0.57 | $<10^{-61}$ |
| Huextetitla 2019 | 250 kb | 0.57 | $<10^{-61}$ |
| *Santa Cruz 2020* | 500 kb | 0.63 | $<10^{-81}$ |
| *Chapulhuacanito 2021* | 500 kb | 0.66 | $<10^{-88}$ |
| Chapulhuacanito 2003 | 500 kb | 0.63 | $<10^{-78}$ |
| Chapulhuacanito 2006 | 500 kb | 0.65 | $<10^{-85}$ |
| Chapulhuacanito 2017 | 500 kb | 0.64 | $<10^{-84}$ |
| Huextetitla 2003 | 500 kb | 0.64 | $<10^{-83}$ |
| Huextetitla 2019 | 500 kb | 0.62 | $<10^{-75}$ |

**Table S5.** Results of the analysis of the partial correlation between local minor parent ancestry and the number of linked coding and conserved basepairs. Since recombination rate has a strong impact on local ancestry (Table S4), this analysis attempts to control for this effect by accounting for local recombination rate. We repeated this analysis in windows of a particular genetic length (Table S6). As before, we thinned windows to include only one per Mb. Population names in italics indicate the focal samples primarily discussed in the main text.

| **Population** | **Window size** | **Partial correlation recombination rate (p-value)** | **Partial correlation coding (p-value)** | **Partial correlation conserved (p-value)** |
|---|---|---|---|---|
| *Santa Cruz 2020* | 100 kb | 0.48 ($<10^{-41}$) | 0.08 (0.039) | -0.03 (0.40) |
| *Chapulhuacanito 2021* | 100 kb | 0.54 ($<10^{-54}$) | 0.02 (0.58) | -0.02 (0.61) |
| Chapulhuacanito 2003 | 100 kb | 0.49 ($<10^{-42}$) | -0.018 (0.64) | -0.009 (0.81) |
| Chapulhuacanito 2006 | 100 kb | 0.50 ($<10^{-45}$) | 0.035 (0.35) | -0.008 (0.83) |
| Chapulhuacanito 2017 | 100 kb | 0.55 ($<10^{-54}$) | -0.028 (0.45) | 0.002 (0.95) |
| Huextetitla 2003 | 100 kb | 0.49 ($<10^{-41}$) | 0.045 (0.19) | -0.017 (0.65) |
| Huextetitla 2019 | 100 kb | 0.48 ($<10^{-41}$) | 0.02 (0.56) | -0.001 (0.96) |
| *Santa Cruz 2020* | 250 kb | 0.57 ($<10^{-62}$) | 0.11 (0.005) | -0.088 (0.019) |
| *Chapulhuacanito 2021* | 250 kb | 0.58 ($<10^{-63}$) | 0.055 (0.14) | -0.078 (0.035) |
| Chapulhuacanito 2003 | 250 kb | 0.56 ($<10^{-59}$) | 0.034 (0.37) | -0.061 (0.11) |
| Chapulhuacanito 2006 | 250 kb | 0.58 ($<10^{-63}$) | 0.056 (0.14) | -0.046 (0.22) |
| Chapulhuacanito 2017 | 250 kb | 0.60 ($<10^{-69}$) | 0.017 (0.66) | -0.064 (0.09) |
| Huextetitla 2003 | 250 kb | 0.55 ($<10^{-57}$) | 0.089 (0.017) | -0.062 (0.09) |
| Huextetitla 2019 | 250 kb | 0.56 ($<10^{-57}$) | 0.068 (0.07) | -0.024 (0.51) |
| *Santa Cruz 2020* | 500 kb | 0.62 ($<10^{-77}$) | 0.04 (0.29) | -0.022 (0.56) |
| *Chapulhuacanito 2021* | 500 kb | 0.65 ($<10^{-85}$) | 0.00002 (0.99) | -0.008 (0.84) |
| Chapulhuacanito 2003 | 500 kb | 0.62 ($<10^{-76}$) | -0.022 (0.56) | -0.022 (0.55) |
| Chapulhuacanito 2006 | 500 kb | 0.64 ($<10^{-82}$) | 0.017 (0.64) | 0.027 (0.47) |
| Chapulhuacanito 2017 | 500 kb | 0.64 ($<10^{-82}$) | -0.04 (0.28) | 0.005 (0.89) |
| Huextetitla 2003 | 500 kb | 0.63 ($<10^{-79}$) | 0.044 (0.24) | -0.011 (0.24) |
| Huextetitla 2019 | 500 kb | 0.61 ($10^{-73}$) | 0.011 (0.77) | -0.011 (0.77) |

**Table S6.** Analysis of correlations between minor parent ancestry and the number of coding or conserved basepairs in a window of a given genetic size. Since recombination rate has a strong effect on minor parent ancestry, and recombination rate is strongly correlated with functional elements in swordtails (see Supporting Information 3), analyses in genetic windows allows us to disentangle these effects. Windows were thinned so that on average on window per Mb was retained, and the same number of windows was retained for each analysis. Population names in italics indicate the focal samples primarily discussed in the main text.

| **Population** | **Window size** | **Correlation coding (p-value)** | **Correlation conserved (p-value)** |
|---|---|---|---|
| *Santa Cruz 2020* | 0.1 cM | -0.15 (0.0002) | -0.18 ($<10^{-8}$) |
| *Chapulhuacanito 2021* | 0.1 cM | -0.21 ($<10^{-7}$) | -0.27 ($<10^{-10}$) |
| Chapulhuacanito 2003 | 0.1 cM | -0.19 ($10^{-5}$) | -0.25 ($<10^{-8}$) |
| Chapulhuacanito 2006 | 0.1 cM | -0.21 ($<10^{-6}$) | -0.27 ($10^{-11}$) |
| Chapulhuacanito 2017 | 0.1 cM | -0.22 ($<10^{-7}$) | -0.29 ($<10^{-11}$) |
| Huextetitla 2003 | 0.1 cM | -0.14 (0.0005) | -0.23 ($<10^{-8}$) |
| Huextetitla 2019 | 0.1 cM | -0.11 (0.006) | -0.19 ($<10^{-5}$) |
| *Santa Cruz 2020* | 0.25 cM | -0.23 ($10^{-7}$) | -0.29 ($<10^{-10}$) |
| *Chapulhuacanito 2021* | 0.25 cM | -0.26 ($<10^{-8}$) | -0.29 ($<10^{-10}$) |
| Chapulhuacanito 2003 | 0.25 cM | -0.26 ($<10^{-8}$) | -0.28 ($<10^{-9}$) |
| Chapulhuacanito 2006 | 0.25 cM | -0.26 ($<10^{-8}$) | -0.30 ($<10^{-11}$) |
| Chapulhuacanito 2017 | 0.25 cM | -0.27 ($<10^{-9}$) | -0.28 ($<10^{-9}$) |
| Huextetitla 2003 | 0.25 cM | -0.22 ($<10^{-6}$) | -0.29 ($<10^{-10}$) |
| Huextetitla 2019 | 0.25 cM | -0.19 (0.00002) | -0.25 ($<10^{-7}$) |
| *Santa Cruz 2020* | 0.5 cM | -0.34 ($<10^{-19}$) | -0.47 ($<10^{-29}$) |
| *Chapulhuacanito 2021* | 0.5 cM | -0.38 ($<10^{-23}$) | -0.42 ($<10^{-29}$) |
| Chapulhuacanito 2003 | 0.5 cM | -0.38 ($<10^{-22}$) | -0.42 ($<10^{-27}$) |
| Chapulhuacanito 2006 | 0.5 cM | -0.38 ($<10^{-24}$) | -0.41 ($<10^{-27}$) |
| Chapulhuacanito 2017 | 0.5 cM | -0.39 ($<10^{-25}$) | -0.42 ($<10^{-28}$) |
| Huextetitla 2003 | 0.5 cM | -0.33 ($<10^{-18}$) | -0.41($<10^{-27}$) |
| Huextetitla 2019 | 0.5 cM | -0.32 ($<10^{-16}$) | -0.38 ($<10^{-23}$) |

**Table S7.** Summary of local ancestry correlations across populations in windows of different sizes. Analysis was conducted on thinned data, so that for each data set one window per Mb was retained, resulting in the same number of windows across comparisons. Both *X. birchmanni* x *X. cortezi* (Santa Cruz, Chapulhuacanito, Huextetitla) and *X. birchmanni* x *X. malinche* populations (Acuapa, Aguazarca, Tlatemaco) are included in this analysis. Note that the *X. birchmanni* x *X. cortezi* populations at Santa Cruz and Huextetitla occur in the same river system. All *X. birchmanni* x *X. malinche* populations occur in different river systems.

| **Population 1** | **Population 2** | **Window size** | **Correlation in minor parent ancestry (Spearman's ρ)** | **P-value** |
|---|---|---|---|---|
| Santa Cruz 2020 | Chapulhuacanito 2021 | 500 kb | 0.86 | $<10^{-100}$ |
| Santa Cruz 2020 | Huextetitla 2019 | 500 kb | 0.92 | $<10^{-100}$ |
| Chapulhuacanito 2021 | Chapulhuacanito 2017 | 500 kb | 0.94 | $<10^{-100}$ |
| Chapulhuacanito 2021 | Acuapa 2018 | 500 kb | 0.20 | $<10^{-7}$ |
| Chapulhuacanito 2021 | Tlatemaco 2017 | 500 kb | 0.13 | 0.0007 |
| Chapulhuacanito 2021 | Aguazarca 2016 | 500 kb | 0.16 | $<10^{-4}$ |
| Santa Cruz 2020 | Chapulhuacanito 2021 | 250 kb | 0.82 | $<10^{-100}$ |
| Santa Cruz 2020 | Huextetitla 2019 | 250 kb | 0.90 | $<10^{-100}$ |
| Chapulhuacanito 2021 | Chapulhuacanito 2017 | 250 kb | 0.92 | $<10^{-100}$ |
| Chapulhuacanito 2021 | Acuapa 2018 | 250 kb | 0.19 | $<10^{-6}$ |
| Chapulhuacanito 2021 | Tlatemaco 2017 | 250 kb | 0.10 | 0.012 |
| Chapulhuacanito 2021 | Aguazarca 2016 | 250 kb | 0.14 | 0.0003 |
| Santa Cruz 2020 | Chapulhuacanito 2021 | 100 kb | 0.79 | $<10^{-100}$ |
| Santa Cruz 2020 | Huextetitla 2019 | 100 kb | 0.87 | $<10^{-100}$ |
| Chapulhuacanito 2021 | Chapulhuacanito 2017 | 100 kb | 0.91 | $<10^{-100}$ |
| Chapulhuacanito 2021 | Acuapa 2018 | 100 kb | 0.19 | $<10^{-5}$ |
| Chapulhuacanito 2021 | Tlatemaco 2017 | 100 kb | 0.05 | 0.20 |
| Chapulhuacanito 2021 | Aguazarca 2016 | 100 kb | 0.14 | 0.0002 |

**Table S8.** Summary of local ancestry correlations across populations in windows of different genetic size. Binning into windows of varying genetic size allows us to partially control for the strong relationship between minor parent ancestry and recombination rate. We performed two analyses using a partial correlation approach, one comparing ancestry across populations while accounting for covariance due to the number of coding basepairs in a window, and a separate analysis comparing ancestry across populations while accounting for covariance due to the number of conserved basepairs in a window. Both *X. birchmanni* x *X. cortezi* (Santa Cruz, Chapulhuacanito, Huextetitla) and *X. birchmanni* x *X. malinche* populations (Acuapa, Aguazarca, Tlatemaco) are included in this analysis. Note that the *X. birchmanni* x *X. cortezi* populations at Santa Cruz and Huextetitla occur in the same river system. All *X. birchmanni* x *X. malinche* populations occur in different river systems. Analysis was conducted on thinned data, so that for each data set one window per 1.5 cMs was retained, resulting in the same number of windows analyzed across comparisons.

| **Population 1** | **Population 2** | **Window size (cM)** | **Partial correlation in minor parent ancestry (p-value)** | **Partial correlation with conserved basepair density (p-value)** |
|---|---|---|---|---|
| Santa Cruz 2020 | Chapulhuacanito 2021 | 0.5 | 0.81 ($<10^{-100}$) | -0.13 (<0.001) |
| Santa Cruz 2020 | Huextetitla 2019 | 0.5 | 0.93 ($<10^{-100}$) | -0.17 ($10^{-5}$) |
| Chapulhuacanito 2021 | Chapulhuacanito 2017 | 0.5 | 0.95 ($<10^{-100}$) | -0.08 (0.036) |
| Chapulhuacanito 2021 | Acuapa 2018 | 0.5 | 0.22 ($<10^{-8}$) | -0.42 ($<10^{-28}$) |
| Chapulhuacanito 2021 | Tlatemaco 2017 | 0.5 | -0.05 (0.21) | -0.44 ($<10^{-31}$) |
| Chapulhuacanito 2021 | Aguazarca 2016 | 0.5 | 0.10 (0.01) | -0.43 ($<10^{-30}$) |
| Santa Cruz 2020 | Chapulhuacanito 2021 | 0.25 | 0.78 ($<10^{-100}$) | -0.08 (0.03) |
| Santa Cruz 2020 | Huextetitla 2019 | 0.25 | 0.93 ($<10^{-100}$) | -0.15 (0.0001) |
| Chapulhuacanito 2021 | Chapulhuacanito 2017 | 0.25 | 0.95 ($<10^{-100}$) | -0.10 (0.016) |
| Chapulhuacanito 2021 | Acuapa 2018 | 0.25 | 0.18 ($<10^{-5}$) | -0.37 ($<10^{-21}$) |
| Chapulhuacanito 2021 | Tlatemaco 2017 | 0.25 | -0.02 (0.44) | -0.39 ($<10^{-23}$) |
| Chapulhuacanito 2021 | Aguazarca 2016 | 0.25 | 0.06 (0.12) | -0.39 ($<10^{-22}$) |
| Santa Cruz 2020 | Chapulhuacanito 2021 | 0.1 | 0.79 ($<10^{-100}$) | -0.02 (0.58) |

| | | | | |
|---|---|---|---|---|
| Santa Cruz 2020 | Huextetitla 2019 | 0.1 | 0.91 ($<10^{-100}$) | -0.21 ($<10^{-7}$) |
| Chapulhuacanito 2021 | Chapulhuacanito 2017 | 0.1 | 0.94 ($<10^{-100}$) | -0.014 (0.71) |
| Chapulhuacanito 2021 | Acuapa 2018 | 0.1 | 0.18 ($<10^{-5}$) | -0.31 ($<10^{-15}$) |
| Chapulhuacanito 2021 | Tlatemaco 2017 | 0.1 | 0.038 (0.34) | -0.32 ($<10^{-16}$) |
| Chapulhuacanito 2021 | Aguazarca 2016 | 0.1 | 0.13 (0.0008) | -0.32 ($<10^{-16}$) |

**Table S9.** Given the high correlations in minor parent ancestry observed between independently formed hybrid populations at Santa Cruz and Chapulhuacanito, we performed a series of analyses to exclude the role of potential artifacts in generating the signal we observe (see Methods). Thinned ancestry informative sites – ancestry informative sites were thinned before the HMM was run to better control for variation in power to infer ancestry across different regions of the genome. Intervals where ancestry informative sites occurred at higher than the median density were thinned to the median density and the HMM was re-run. Low power windows removed – windows with the fewest ancestry informative sites (lowest 5% quantile) were removed before analysis. Chromosome ends removed – windows within the last Mb of chromosomes, some of which overlapped with telomeric sequences, were removed. Repeats excluded – ancestry informative sites that overlapped with annotated repeats were excluded before analysis. All analyses were conducted on spatially thinned windows, so that for each data set one window per Mb was retained, resulting in the same number of windows across comparisons.

| **Population 1** | **Population 2** | **Method** | **Window size** | **Spearman's ρ (p-value)** |
|---|---|---|---|---|
| Santa Cruz 2020 | Chapulhuacanito 2021 | Thinned ancestry informative sites | 500 kb | 0.86 ($<10^{-100}$) |
| Santa Cruz 2020 | Chapulhuacanito 2021 | Thinned ancestry informative sites | 250 kb | 0.82 ($<10^{-100}$) |
| Santa Cruz 2020 | Chapulhuacanito 2021 | Thinned ancestry informative sites | 100 kb | 0.79 ($<10^{-100}$) |
| Santa Cruz 2020 | Chapulhuacanito 2021 | Population specific priors | 500 kb | 0.87 ($<10^{-100}$) |
| Santa Cruz 2020 | Chapulhuacanito 2021 | Population specific priors | 250 kb | 0.84 ($<10^{-100}$) |
| Santa Cruz 2020 | Chapulhuacanito 2021 | Population specific priors | 100 kb | 0.80 ($<10^{-100}$) |
| Santa Cruz 2020 | Chapulhuacanito 2021 | Low power windows removed | 500 kb | 0.85 ($<10^{-100}$) |
| Santa Cruz 2020 | Chapulhuacanito 2021 | Low power windows removed | 250 kb | 0.82 ($<10^{-100}$) |
| Santa Cruz 2020 | Chapulhuacanito 2021 | Low power windows removed | 100 kb | 0.79 ($<10^{-100}$) |
| Santa Cruz 2020 | Chapulhuacanito 2021 | Chromosome ends removed | 500 kb | 0.87 ($<10^{-100}$) |
| Santa Cruz 2020 | Chapulhuacanito 2021 | Chromosome ends removed | 250 kb | 0.84 ($<10^{-100}$) |
| Santa Cruz 2020 | Chapulhuacanito 2021 | Chromosome ends removed | 100 kb | 0.80 ($<10^{-100}$) |
| Santa Cruz 2020 | Chapulhuacanito 2021 | Repeats excluded | 500 kb | 0.86 ($<10^{-100}$) |

| | | | | |
|---|---|---|---|---|
| Santa Cruz 2020 | Chapulhuacanito 2021 | Repeats excluded | 250 kb | 0.83 ($<10^{-100}$) |
| Santa Cruz 2020 | Chapulhuacanito 2021 | Repeats excluded | 100 kb | 0.79 ($<10^{-20}$) |
| Santa Cruz 2020 | Chapulhuacanito 2021 | Low power removed and chromosome ends removed | 500 kb | 0.87 ($<10^{-100}$) |
| Santa Cruz 2020 | Chapulhuacanito 2021 | Low power removed and chromosome ends removed | 250 kb | 0.84 ($<10^{-100}$) |
| Santa Cruz 2020 | Chapulhuacanito 2021 | Low power removed and chromosome ends removed | 100 kb | 0.80 ($<10^{-100}$) |

*Attached excel table:* **Table S10.** Genes found in shared minor parent islands. Bolded rows indicate islands that changed significantly in ancestry over time at $p<0.05$.

**Table S11.** Identity of mitochondrial haplotypes found in both *X. birchmanni* x *X. cortezi* populations and in time series data from Chapulhuacanito.

| **Population** | **Number of *X. cortezi* mitochondrial haplotypes** | **Number of *X. birchmanni* mitochondrial haplotypes** | **Frequency *X. cortezi* haplotype** |
|---|---|---|---|
| Santa Cruz 2020 | 242 | 0 | 100% |
| Chapulhuacanito 2003 | 7 | 0 | 100% |
| Chapulhuacanito 2006 | 19 | 0 | 100% |
| Chapulhuacanito 2017 | 18 | 1 | 95% |
| Chapulhuacanito 2021 | 69 | 0 | 100% |

**Table S12.** Data from *Xiphophorus* species that was used to infer ancestral sequence states for LDhelmet.

| **Species** | **Data source** | **SRA accession** |
|---|---|---|
| *X. malinche* | Schumer et al. 2018 | SRX3626998-SRX3626999 |
| *X. birchmanni* | Schumer et al. 2018 | SRX2486382-SRX3648154 |
| *X. cortezi* | Powell et al. 2020 | SRX7860174-SRX7860180 |
| *X. montezumae* | Schumer et al. 2016 | SRR3086791 |
| *X. continens* | Preising et al. 2022 | NA |
| *X. nezahualcoyotl* | Schumer et al. 2016 | SRR3086878; SRR3086631 |
| *X. nigrensis* | Preising et al. 2022 | NA |
| *X. multilineatus* | Preising et al. 2022 | NA |
| *X. variatus* | Powell et al. 2020 | SRR13982900 |
| *X. maculatus* | Schartl et al. 2013 | SRR7525605 |
| *X. hellerii* | Shen et al. 2016 | SRR7532852 |

**Table S13.** Summary statistics input for ABCreg demographic inference analysis of Chapulhuacanito and Santa Cruz populations.

| **Whole genome** | | | | |
|---|---|---|---|---|
| *Population* | *Median minor parent tract length* | *Average ancestry (proportion X. cortezi*) | *Standard deviation in genome-wide ancestry* | *Coefficient of variation in ancestry* |
| Chapulhuacanito 2021 | 69.6 kb | 0.76 | 0.018 | 0.024 |
| Santa Cruz 2020 | 33.9 kb | 0.85 | 0.031 | 0.036 |
| **Chromosome 2** | | | | |
| Chapulhuacanito 2021 | 63.6 kb | 0.80 | 0.053 | 0.067 |
| Santa Cruz 2020 | 30.0 kb | 0.86 | 0.067 | 0.078 |

*Attached excel document:* **Table S14.** Significantly enriched gene otology categories identified among genes in minor parent islands.

### Supporting Information References


1. Li H. Minimap2: pairwise alignment for nucleotide sequences. Bioinformatics. 2018;34: 3094–3100. doi:10.1093/bioinformatics/bty191

2. Schumer M, Xu C, Powell DL, Durvasula A, Skov L, Holland C, et al. Natural selection interacts with recombination to shape the evolution of hybrid genomes. Science. 2018;360: 656. doi:10.1126/science.aar3684

3. Baker Z, Schumer M, Haba Y, Bashkirova L, Holland C, Rosenthal GG, et al. Repeated losses of PRDM9-directed recombination despite the conservation of PRDM9 across vertebrates. In: eLife [Internet]. 6 Jun 2017 [cited 23 Jul 2019]. doi:10.7554/eLife.24133

4. Cavassim MIA, Baker Z, Hoge C, Schierup MH, Schumer M, Przeworski M. PRDM9 losses in vertebrates are coupled to those of paralogs ZCWPW1 and ZCWPW2. bioRxiv; 2021. p. 2021.06.08.447603. doi:10.1101/2021.06.08.447603

5. Singhal S, Leffler EM, Sannareddy K, Turner I, Venn O, Hooper DM, et al. Stable recombination hotspots in birds. Science. 2015;350: 928–932. doi:10.1126/science.aad0843

6. Lam I, Keeney S. Nonparadoxical evolutionary stability of the recombination initiation landscape in yeast. Science. 2015;350: 932–937. doi:10.1126/science.aad0814

7. Armstrong J, Hickey G, Diekhans M, Deran A, Fang Q, Xie D, et al. Progressive alignment with Cactus: a multiple-genome aligner for the thousand-genome era. bioRxiv. 2019; 730531. doi:10.1101/730531

8. Powell DL, Moran BM, Kim BY, Banerjee SM, Aguillon SM, Fascinetto-Zago P, et al. Two new hybrid populations expand the swordtail hybridization model system. Evolution. 2021;75: 2524–2539. doi:10.1111/evo.14337

9. Moran BM, Payne C, Langdon Q, Powell DL, Brandvain Y, Schumer M. The genomic consequences of hybridization. Wittkopp PJ, editor. eLife. 2021;10: e69016. doi:10.7554/eLife.69016

10. Langdon QK, Powell DL, Kim B, Banerjee SM, Payne C, Dodge TO, et al. Predictability and parallelism in the contemporary evolution of hybrid genomes. PLOS Genetics. 2022;18: e1009914. doi:10.1371/journal.pgen.1009914

11. Cui R, Schumer M, Rosenthal GG. Admix'em: a flexible framework for forward-time simulations of hybrid populations with selection and mate choice. Bioinformatics. 2016;32: 1103–1105. doi:10.1093/bioinformatics/btv700

12. Harris K, Nielsen R. The Genetic Cost of Neanderthal Introgression. Genetics. 2016;203: 881–891. doi:10.1534/genetics.116.186890

13. Matute DR, Comeault AA, Earley E, Serrato-Capuchina A, Peede D, Monroy-Eklund A, et al. Rapid and Predictable Evolution of Admixed Populations Between Two Drosophila Species Pairs. Genetics. 2019 [cited 20 Apr 2020]. doi:10.1534/genetics.119.302685

14. Veller C, Edelman NB, Muralidhar P, Nowak MA. Recombination and Selection Against Introgressed DNA. Evolution. 2023; qpad021. doi:10.1093/evolut/qpad021

15. Schumer M, Powell DL, Corbett-Detig R. Versatile simulations of admixture and accurate local ancestry inference with mixnmatch and ancestryinfer. Mol Ecol Resour. 2020;20: 1141–1151. doi:10.1111/1755-0998.13175

16. Gravel S. Population Genetics Models of Local Ancestry. Genetics. 2012;191: 607. doi:10.1534/genetics.112.139808

17. Durinck S, Spellman PT, Birney E, Huber W. Mapping identifiers for the integration of genomic datasets with the R/Bioconductor package biomaRt. Nat Protoc. 2009;4: 1184–1191. doi:10.1038/nprot.2009.97

18. Falcon S, Gentleman R. Using GOstats to test gene lists for GO term association. Bioinformatics. 2007;23: 257–258. doi:10.1093/bioinformatics/btl567

19. Morgan, Martin, Falcon S, Gentleman R. GSEABase: Gene set enrichment data structures and methods version 1.52.1 from Bioconductor. [cited 5 Aug 2021]. Available: https://rdrr.io/bioc/GSEABase/

20. Moran BM, Payne CY, Powell DL, Iverson ENK, Banerjee SM, Langdon QK, et al. A Lethal Genetic Incompatibility between Naturally Hybridizing Species in Mitochondrial Complex I. 2021 Jul p. 2021.07.13.452279. doi:10.1101/2021.07.13.452279

21. Drew K, Lee C, Huizar RL, Tu F, Borgeson B, McWhite CD, et al. Integration of over 9,000 mass spectrometry experiments builds a global map of human protein complexes. Molecular Systems Biology. 2017;13: 932. doi:10.15252/msb.20167490

22. Drost H-G, Gabel A, Grosse I, Quint M. Evidence for Active Maintenance of Phylotranscriptomic Hourglass Patterns in Animal and Plant Embryogenesis. Molecular Biology and Evolution. 2015;32: 1221–1231. doi:10.1093/molbev/msv012

23. Groh JS, Coop G. The temporal and genomic scale of selection following hybridization. bioRxiv; 2023. p. 2023.05.25.542345. doi:10.1101/2023.05.25.542345
